# A dual-receptor model of serotonergic psychedelics

**DOI:** 10.1101/2024.04.12.589282

**Authors:** Arthur Juliani, Veronica Chelu, Laura Graesser, Adam Safron

## Abstract

Serotonergic psychedelics have been identified as promising next-generation therapeutic agents in the treatment of mood and anxiety disorders. While their efficacy has been increasingly validated, the mechanism by which they exert a therapeutic effect is still debated. A popular theoretical account is that excessive 5-HT2a agonism disrupts cortical dynamics, relaxing the precision of maladaptive high-level beliefs and making them more malleable and open to revision. We extend this perspective by developing a simple energy-based model of cortical dynamics based on predictive processing which incorporates effects of neuromodulation. Using this model, we propose and simulate hypothetical computational mechanisms for both 5-HT2a and 5-HT1a agonism. Results from our model are able to account for a number of existing empirical observations concerning serotonergic psychedelics effects on cognition and affect. Using the findings of our model, we provide a theoretically-grounded hypothesis for the clinical success of LSD, psilocybin, and DMT, as well as identify the design space of biased 5-HT1a agonist psychedelics such as 5-MeO-DMT as potentially fruitful in the development of more effective and tolerable psychotherapeutic agents in the future.

## 1 Introduction

Serotonergic psychedelics have received significant attention from both research scientists and clinicians in recent years for their potential to treat a variety of psychiatric conditions ranging from depression and anxiety to substance use and obsessive compulsive disorders (M. W. Johnson, Hendricks, Barrett, & Griffiths, 2019; Sessa, 2018; Griffiths et al., 2016; M. W. Johnson & Griffiths, 2017; Carhart-Harris et al., 2021; Timmermann, Zeifman, Erritzoe, Nutt, & Carhart-Harris, 2024). This transdiagnostic efficacy has led some researchers to hypothesize that there may be a single primary underlying factor of psychopathology which psychedelic therapy is acting to address (Carhart-Harris et al., 2022). Despite significant progress in understanding the empirical therapeutic effects of these drugs, theoretical models which describe and predict these effects are less well developed. While the key role of the 5-HT2a receptor system and downstream neurotrophic effects are largely accepted by the scientific community (Shao et al., 2021; Calder & Hasler, 2023; Banushi & Polito, 2023), the role of subjective experience in the therapeutic effects of psychedelics is still a topic of considerable debate (Yaden & Griffiths, 2020; Cameron et al., 2021).

A variety of competing theories have been developed to explain the mechanics behind the subjective effects of psychedelics and their relationship to therapeutic outcomes (Doss et al., 2022). Among these, the *Relaxed Beliefs Under Psychedelics* (REBUS) model (Carhart-Harris & Friston, 2019) has received additional attention for its translational application in guiding clinical protocol development. For example, it is compatible with “neural annealing theory,” which has gained popularity among practitioners of psychedelic-assisted therapy as an explanatory framework in recent years (M. Johnson, 2019; Gómez-Emilsson, 2021). According to the REBUS model and neural annealing theory, serotonergic psychedelics exert their therapeutic effect by relaxing the precision of high-level beliefs both acutely and post-acutely, thus making them more amenable to modification through introspection and interpersonal therapy than would otherwise be possible. This hypothesis is supported by evidence of an increase in the entropy of cortical transition dynamics during the acute phase of psychedelic use (Singleton et al., 2022), and a relaxation of long-term propositional beliefs after psychedelic therapy (Zeifman et al., 2022).

Despite some preliminary evidence in support of REBUS, it is not clear whether we should expect the relaxation of beliefs to hold true across the dose-response curve for all substances in the broad class of serotonergic psychedelics. One piece of evidence which would suggest that serotonergic psychedelics may transiently strengthen beliefs in some cases are frequent reports of insight-experiences (Laukkonen et al., 2023; McGovern et al., 2023), in which individuals arrive a strongly-felt (but potentially erroneous) new beliefs during the acute drug experience. Another is the experience of pareidolia, in which ambiguous low-level visual stimuli are perceived as belonging to a higher-level precept, such as seeing a face in a cloud (Liu et al., 2014; Lhotka, Ischebeck, Helmlinger, & Zaretskaya, 2023). This is most extreme in cases of so-called “entities” and “alternate realities” under the effects of DMT, which are often described as appearing more real than normal conscious experience (Luan et al., 2023).

The recently introduced *Altered Beliefs Under Psychedelics* (ALBUS) model was developed to attempt to account for this wider set of potential effects on belief representations, with the therapeutic belief relaxation seen in clinical studies being a special case which results from specific dose ranges and controlled environmental influences (Safron, 2020). While this alternative hypothesis is based on a study of the theoretical non-linear dynamics in the underlying activity of the relevant circuits in the cortex, it currently lacks empirical validation and is under-specified in a number of ways which limit its practical application as a predictive tool.

Although the purported mechanism behind the disruption of belief representation is 5-HT2a agonism, there is also a non-trivial contribution by other serotonin (5-HT) receptor populations in the changes to neural dynamics which result from serotonergic psychedelics, with activity at 5-HT1a receptors in particular exerting a significant effect (Deco et al., 2018; Singleton et al., 2022). Psilocybin, LSD, and DMT all have significant affinity for the 5-HT1a receptor in addition to the 5-HT2a receptor (Rickli, Moning, Hoener, & Liechti, 2016). Rather than serving a minor or even completely negligible role, as has been previously assumed, we first provide a review of empirical evidence which provide insight into the key role 5-HT1a agonism may play in psychedelic experiences and therapeutic outcomes.

We then present a computational model inspired by predictive processing (PP) which incorporates hypothesized mechanisms of 5-HT2a and 5-HT1a agonism on the process of belief representation in the cortex. Results from our model are consistent with empirical findings that although 5-HT2a agonism is necessary and sufficient for psychedelic phenomenology, 5-HT1a agonism may play a significant role in modulating the acute experience and clinical outcomes. Based on these findings, we end with a discussion of the potential implications of our model for clinical research and psychedelic drug development. Properly characterizing the modulatory role played by 5-HT1a agonism in the psychedelic experience may be essential to the development of next-generation psychedelic substances which provide both greater psychological tolerability as well as more consistent long-term therapeutic effects (Warren et al., 2024).

### 1.1 Serotonergic neuromodulation as a stress response system

One view of the serotonin system is that it serves to mediate cognitive and affective responses to stress (Puglisi-Allegra & Andolina, 2015; Murnane, 2019; Zhong, Li, Feng, & Luo, 2017; Hayashi, Nakao, & Nakamura, 2015). This can be seen as an extension of a broader evolutionarily preserved role for serotonin in responding to aversive stimuli (Dayan & Huys, 2009). Multiple models within this framework view serotonin signaling as composed of two unique interdependent systems, corresponding to the two main serotonin receptor populations in the brain. In one model, these receptor systems instantiate separate “active” and “passive” coping strategies which an organism may engage under stress (Carhart-Harris & Nutt, 2017). In another, they been identified as “flux” and “automatic” cognitive states (Shine et al., 2022), which, while not identical, can be understood to broadly perform similar roles to the two coping strategies described by Carhart-Harris and Nutt.

The human nervous system contains over a dozen unique serotonin receptors (Nichols & Nichols, 2008). Of these, 5-HT1a and 5-HT2a are both the most widely distributed within the brain and the most well studied (Beliveau et al., 2017). Of particular interest are the role of the postsynaptically expressed 5-HT1a and 5-HT2a receptors in the cortex. These receptor populations are understood to operate in relatively simple opponency with one another within cortical pyramidal cells, with 5-HT1a receptors inhibiting and 5-HT2a receptors exciting the postsynaptic cell (Aghajanian & Marek, 1997; Puig & Gulledge, 2011). The two stress response systems within the brain are hypothesized to be instantiated by 5-HT2a and 5-HT1a receptor systems (Carhart-Harris & Nutt, 2017; Shine et al., 2022).

Within this dual-strategy framework, small to moderate amounts of stress activate the 5-HT1a mediated passive coping system while large amounts of stress activate the 5-HT2a mediated active coping system. In passive coping, the animal maintains the current behavioral policy but modulates affect in response to manageable levels of stress. In contrast, active coping is engaged when the level of stress is significant enough to require the instantiation of novel and divergent internal models, beliefs, or behavioral policies (Brouwer & Carhart-Harris, 2021; Safron & Sheikhbahaee, 2021). This bi-modal behavioral distribution can be seen in a simplified form in serotonin-mediated rodent responses to stress from predatory threats, where a low-level threat initiates freezing behavior while a high-level threat initiates fleeing behavior (Seo et al., 2019).

Importantly, both coping strategies are mediated by the same system of serotonergic neuromodulation. Within a stress-response model of serotonin function, the extent to which a given strategy is deployed is a function of 5-HT release in the dorsal raphae nucleus (DRN). A complete account of the computational role of the DRN is still being developed, but evidence exists that DRN neurons may compute unsigned prediction errors (Matias, Lottem, Dugué, & Mainen, 2017), and that the magnitude of this signal corresponds to downstream 5-HT release in the cortex. This prediction error computation is hypothesized to be part of a larger role for the DRN in value prediction complementary to that of the ventral tegmental area (Harkin, Grossman, Cohen, Béïque, & Naud, 2023). The response of the serotonin system to stress may therefore be mediated by the ability of the organism to form and maintain successfully predictive representations.

Within this framework, 5-HT neuromodulation then serves a role in responding to stress by improving the computational efficiency of the processes required to learn and maintain accurate and predictive representations. The ability to efficiently learn and maintain adaptive predictive representations is precisely the construct of cognitive flexibility (Ionescu, 2012)—the characteristic that enables animals or humans to adaptively generate appropriate behavioral responses based on changing sensory stimuli—which psychedelics have been demonstrated to improve (Doss et al., 2021; Torrado Pacheco, Olson, Garza, & Moghaddam, 2023), and is believed to be generally associated with serotonin signalling (Matias et al., 2017).

Given that all classic psychedelics have significant affinity for the 5-HT2a receptor system, they have been characterized as a prototypical example of a substance that induces an “active coping” response in an organism by Carhart-Harris and Nutt. This manifests in the acute psychedelic state which enables a state of “metaplasticity”—the dynamic regulation of the extent to which synaptic plasticity can be induced (Abraham & Bear, 1996; Nardou et al., 2023), enabling a re-evaluation of previously held beliefs and the spontaneous assumption of novel representations of self or environment (for a review of this effect across various forms of belief, see (Letheby, 2021)). In contrast, Carhart-Harris and Nutt propose MDMA as an exemplar “passive coping” substance due to its role in releasing endogenous serotonin, which preferentially activates the 5-HT1a receptor system. In humans this is hypothesized to manifest most acutely in the felt sense of equanimity and acceptance that characterizes the unique phenomenology of the MDMA (Greer & Tolbert, 1990; Carhart-Harris & Nutt, 2017).

A complication to this straightforward account is that psilocybin, LSD, and DMT all bind with significant affinity to 5-HT1a receptors as well 5-HT2a receptors (Rickli et al., 2016) (See Figure 1). Given the strong positive correlation between receptor binding affinity and efficacy for this class of drugs (Halberstadt, Chatha, Klein, Wallach, & Brandt, 2020), one would expect them to engage both coping systems simultaneously, as opposed to only engaging the active coping system. Below we present a review of existing evidence that this may be the case.

**Figure 1.**
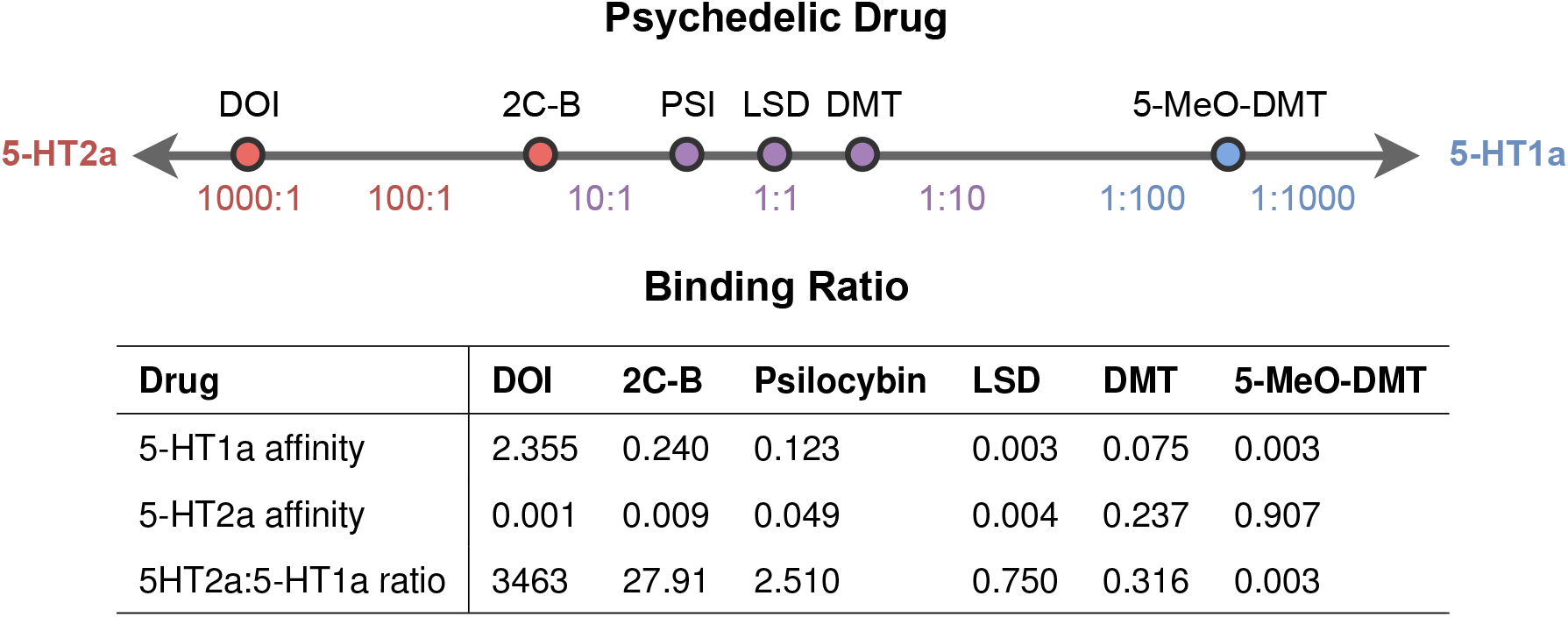
Relative 5-HT1a and 5-HT2a binding affinities *K*_*i*_(*μM*) for select psychedelic substances. Lower values correspond to higher binding affinity. DOI has the highest relative 5-HT2a binding affinity, while 5-MeO-DMT has the highest relative 5-HT1a binding affinity. Data sourced from (Halberstadt et al., 2012; Rickli et al., 2015, 2016).

### 1.2 5-HT2a and 5-HT1a receptor agonism in psychedelics

5-HT2a agonism has been demonstrated to be necessary and sufficient for inducing the “head twitch response” in rodents (Halberstadt, Koedood, Powell, & Geyer, 2011), and this effect is highly correlated with the existence of psychedelic phenomenology in humans (Halberstadt et al., 2020). There is additional evidence however that agonism at the 5-HT1a receptor is responsible for significantly modulating the 5-HT2a mediated effects of serotonergic psychedelics. Here we consider the evidence for this modulating effect across a number of dimensions including behavioral, neural, clinical, and phenomenological.

On the phenomenological level, 5-HT1a agonism has been identified as being the primary driver of stimulus control in rodents trained to discriminate 5-MeO-DMT (Winter, Filipink, Timineri, Helsley, & Rabin, 2000), and there is evidence that it also plays a significant role in the discrimination of LSD and DMT (Reissig, Eckler, Rabin, & Winter, 2005), though not psilocybin (Winter, Rice, Amorosi, & Rabin, 2007) or DOB (a strongly biased 5-HT2a agonist, similar to DOI) (Benneyworth, Smith, Barrett, & Sanders-Bush, 2005). The role of 5-HT1a agonism in stimulus control in these examples is notably correlated with the relative 5-HT2a and 5-HT1a receptor affinities of each drug (Rickli et al., 2016) (see Figure 1), suggesting an underlying pattern whereby psychedelics with greater relative 5-HT1a agonism also manifest phenomenological effects which are more dependent on that agonism.

Despite the fact that the 5-HT1a agonism of psilocybin does not induce stimulus control, there is still evidence for the role it plays in the drug’s behavioral effects, specifically with respect to the modulation of attentional control (Carter et al., 2005), compulsive behavior (Singh et al., 2023), and exploration (Halberstadt et al., 2011). The later study demonstrated that 5-HT1a agonism is likewise primarily implicated in the behavioral and phenomenological effects of 5-MeO-DMT, which is unsurprising, given the high ratio of 5-HT1a to 5-HT2a affinity in this substance.

The behavioral effects in rodents treated with 5-MeO-DMT are accompanied by changes in cortical dynamics in both PFC and visual cortex, both of which are mediated primarily by 5-HT1a agonism (Riga, Soria, Tudela, Artigas, & Celada, 2014; Riga, Bortolozzi, Campa, Artigas, & Celada, 2016; Riga, Lladó-Pelfort, Artigas, & Celada, 2018). One consistent marker of these disruptions is a decrease in fMRI blood-oxygen-level-dependent (BOLD) signal, which is consistent with the inhibitory role of postsynaptic 5-HT1a agonism, particularly in the PFC. 5-HT1a receptor binding maps are also significantly more predictive than any other serotonin binding map except for 5-HT2a in anticipating the changes to brain activity under LSD (Deco et al., 2018), as well as for psilocybin (Singleton et al., 2022).

Importantly, there is also evidence that 5-HT2a agonism may not play an essential role in the antidepressant effects of psychedelics, even independent of its effects on neuroplasticity. DOI, a highly biased 5-HT2a agonists, has recently failed to produce robust therapeutic effects in a rodent model of depression (Qu, Chang, Ma, Wan, & Hashimoto, 2023). There is also mixed evidence for the efficacy of DOI as a potential treatment of anxiety disorders, with a number of studies finding an anxiolytic effect (Dhonnchadha, Hascoët, Jolliet, & Bourin, 2003; Massé, Hascoët, Dailly, & Bourin, 2006), while others finding an anxiogenic effect in rodent models (Masse, Petit-Demouliere, Dubois, Hascoët, & Bourin, 2008; De Paula, Torricelli, Lopreato, Nascimento, & Viana, 2012). In contrast, it has been demonstrated that psilocybin is capable of producing antidepressant effects in rodent models even when the acute effects of 5-HT2a agonism are partially blocked (Hesselgrave, Troppoli, Wulff, Cole, & Thompson, 2021; Shao et al., 2021), potentially implicating 5-HT1a agonism instead in these effects. It has also been demonstrated that the ability of DMT to induce fear-extinction in rodents is mediated by both 5-HT2a and 5-HT1a mechanisms (Werle et al., 2024).

Another proposed mediator of the antidepressant effects of psychedelics is a post-acute increase in cognitive flexibility (Doss et al., 2021). A review of the literature suggests that cognitive flexibility is more frequently associated with preferential 5-HT1a agonism rather than 5-HT2a agonism (Depoortère et al., 2010; Alvarez, Morales, & Amodeo, 2021; Odland, Kristensen, & Andreasen, 2021). Recent evidence also suggests that blocking 5-HT1a agonism in psilocybin impairs the drug’s positive effects on cognitive flexibility in a rodent model of anorexia nervosa (Conn et al., 2023), suggesting an important mediating role for 5-HT1a.

Finally, serotonin itself demonstrates a significantly biased agonism for 5-HT1a over 5-HT2a receptors (Barnes & Sharp, 1999), and there are commonalities between the phenomenological effects of serotonin releasing drugs and 5-HT1a agonists. In a study of the effects of the serotonin releasing drug MDMA, 5-HT2a agonism was found to play only a negligible role in the acute phenomenology (Liechti, Saur, Gamma, Hell, & Vollenweider, 2000), suggesting in contrast a more prominent role of 5-HT1a agonism in the subjective effects of the drug. There is also a research literature on the anxiolytic effects of biased 5-HT1a agonists, such as 8-OH-DPAT, in rodent models of psychopathology (Luscombe, Martin, Hutchins, Gosden, & Heal, 1993; Schreiber & De Vry, 1993), and these effects are consistent with those of MDMA and other 5-HT releasing drugs. Reductions in anxiety and avoidance are key mediators in a variety of common psychopathologies and their treatment by psychedelic therapy (Zeifman, Wagner, Monson, & Carhart-Harris, 2023), suggesting a potential link between these effects and 5-HT1a agonism. Notably, outcomes from MDMA assisted therapy are characterized by a significant acute decrease in experiential avoidance (Mitchell et al., 2023).

Taken together, it is clear that at the very least 5-HT1a agonism plays a non-trivial modulating role in the acute phenomenology and long-term efficacy of serotonergic psychedelics. The question that then arises is in what way 5-HT1a agonism may contribute to these effects when the most apparent changes in subjective experience are mediated by 5-HT2a agonism (Preller et al., 2017). The predictions of a theoretical model of these drug effects should be able to account for both the anxiolytic, sedating, and prosocial effects (Luscombe et al., 1993; Thompson, Callaghan, Hunt, Cornish, & McGregor, 2007) of 5-HT1a agonism as encapsulated in the construct of “passive coping” as well as the stimulating, pareidolic, and insight-inducing effects of 5-HT2a agonism as encapsulated in the construct of “active coping”, both of which we view as necessary to fully characterize the serotonergic psychedelic experience in humans and other animals (M. W. Johnson et al., 2019; Braun et al., 2024).

### 1.3 Predictive processing and psychopathology

The REBUS model is rooted in the idea that the brain uses an internal model of the world to predict sensory input based on movement and past sensory experience (Carhart-Harris & Friston, 2019). This family of theories, collectively referred to as the predictive processing (PP) framework (Clark, 2013, 2016; Keller & Mrsic-Flogel, 2018; Kanai, Komura, Shipp, & Friston, 2015) rely on the idea that an error signal between the predicted and actual sensory input is used to update an internal representation of the world. Within the Bayesian brain perspective in particular (Knill & Pouget, 2004; Doya, Ishii, Pouget, & Rao, 2006), these internal representations are sometimes referred to as beliefs, so we use this term interchangeably with neural response or neural state. PP is often described as a processing hierarchy, and referred to as hierarchical predictive processing (HPP) in the context of the human cortex, whereby higher levels of this hierarchy of functional brain areas send top-down signals to a lower areas in the form of predictions of the bottom-up stimulus input to that area (see Figure 2 for an illustration of a representational layer in the PP model).

**Figure 2.**
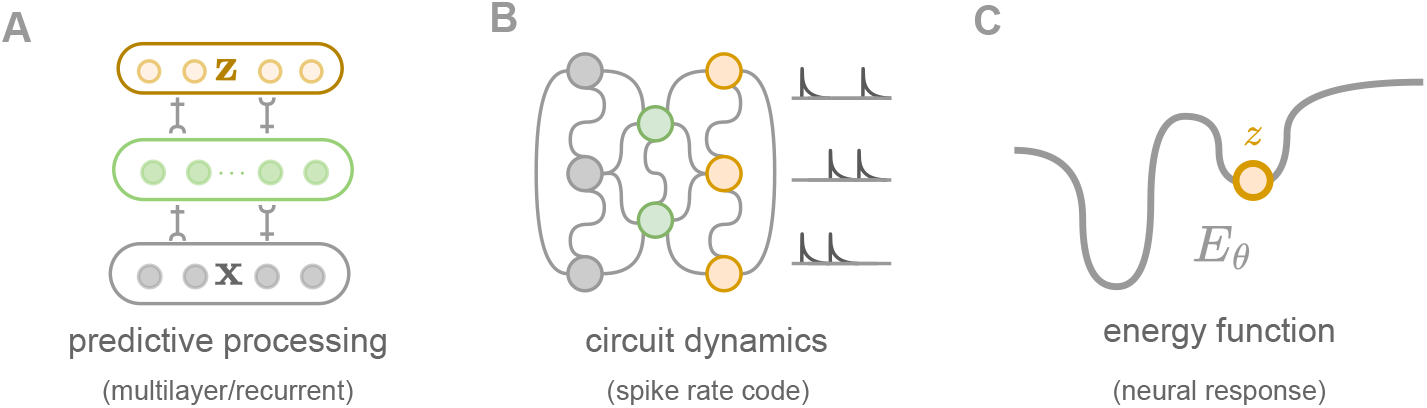
**(A) Theoretical model**. The synaptic connections between the latent neural response vector **z** and the input stimulus **x** is updated based on prediction errors between top-down predictions and stimuli. Prediction errors are sent forward in the hierarchy, while predictions are sent backward. **(B) Population dynamics in the PFC**. Illustrative representation of neural population dynamics in the PFC and associated rate spike code. **(C) Energy function**. A prototypical representation of a 1D energy function induced by the underlying synaptic connectivity structure, along with a sampled neural response *z*.

Within this theoretical framework, it has been proposed prediction errors are dynamically adjusted using a gain modulation mechanism which gates neural plasticity in a context-dependent manner (Behrens, Woolrich, Walton, & Rushworth, 2007; Wilmes & Clopath, 2023; Sadeh, Clopath, Palmer, Frank, & Ziv, 2022). Specifically, the neuromodulatory tone may alter the precision or sensitivity by which cortical circuits prioritize and respond to sensory information or report prediction errors. Representations encoded with greater precision are thus able to exert a greater influence on both upstream and downstream computation. According to the REBUS model, the spontaneous activation of pyramidal neurons due to increased postsynaptic 5-HT2a agonism results in a desynchronization of those neuronal populations, and thus a disruption in their normal ability to robustly encode high-level beliefs or internal representations. It has been speculated, that this disruption leads to a system that is less constrained by prior high-level knowledge, and is thus able to more easily change through experience-dependent learning. Under the REBUS formulation, some psychopathology can be interpreted as maladaptive learning and inference in systems constrained by strong, rigid priors (Carhart-Harris et al., 2022; Juliani, Safron, & Kanai, 2024).

At the neural level, the disruption of high-level functional connectivity induced by psychedelics is accompanied by a decrease in activity and correlation within and between these networks, particularly those involved in self-referential processing (Siegel et al., 2024). This includes regions of the default-mode network (DMN), a brain network that shows increased endogenous fluctuations when an individual is awake but not engaged in a task requiring attention (Kometer, Pokorny, Seifritz, & Volleinweider, 2015; M. W. Johnson et al., 2019), a reconfiguration of communication in the brain characterized by increased brain activity, increased diversity of neural activation patterns compared to normal waking consciousness, elevated Shannon-Boltzmann entropy (Shannon, 1948) of intra-brain-network synchrony (Carhart-Harris et al., 2014), induced “metaplasticity”—thought to be one of the mechanisms underlying the establishment of critical periods (Abraham & Bear, 1996; Nardou et al., 2023).

An implication of connecting neural responses with behavioral fitness is that stable neural responses are not inherently pathological if they are both relevant to the current environmental context and adaptable to future contexts (Juliani et al., 2024). Indeed, such stable, and optimal representations may provide a buffer against various forms of psychopathology (DeYoung & Krueger, 2018). The inverse is also true: even if certain encoded predictions are amenable to change (more adaptable), they can still be maladaptive if they are unable to generalize and instantiate an optimal behavioral policy to ensure success in a given environmental context. Crucially, the ideal situation involves learning models which are both adapted to the current environmental context and malleable enough to make possible future adaptation when the environmental context changes (Mermillod, Bugaiska, & Bonin, 2013).

### 1.4 Prefrontal cortex as an energy-based model

The PFC is involved in higher cognitive functions such as decision-making, working memory, and executive control (Friedman & Robbins, 2022). It is believed to encode the high-level representations of an organism concerning self, others, and world (Miller, Freedman, & Wallis, 2002), as well as the high-level behavioral and attentional policies required to achieve those goals (Domenech & Koechlin, 2015). It is also implicated in a variety of psychopathology (Callicott et al., 2000; Pizzagalli & Roberts, 2022), making it an ideal target region for a pharmacological intervention. Indeed, there is clear neuroimaging evidence that this region is heavily involved in the action of serotonergic psychedelics (Wood, Kim, & Moghaddam, 2012; Luppi et al., 2024; Siegel et al., 2024), with a large density of both 5-HT2a and 5-HT1a postsynaptic receptors and is a primary target of 5-HT neuromodulation arising from the dorsal raphae nucleus (Amargós-Bosch et al., 2004; Puig & Gulledge, 2011). In this work, we propose a mechanism of action of serotonergic psychedelic that uses an abstract and simplified model of neural responses encoded by the PFC.

As a first step, we take a dynamical systems approach, which describes neural population responses as time-varying trajectories in a high-dimensional state space and views the dynamics as acting to shape these trajectories (Khona & Fiete, 2022). A neural manifold is a low-dimensional geometric representation of neural population activity that emerges from high-dimensional neural responses (Langdon, Genkin, & Engel, 2023). It captures the patterns of activity that neurons exhibit during specific tasks or behaviors, summarizing the complex and often heterogeneous activity of individual neurons into a more interpretable form. The manifold represents the constrained set of activity patterns that the population typically explores.

The PFC expresses strong recurrent dynamics (Mante, Sussillo, Shenoy, & Newsome, 2013; J. X. Wang et al., 2018; S. Wang, Falcone, Richmond, & Averbeck, 2023), which are necessary for the formation and maintenance of attractors responsible for the stable responses (Mante et al., 2013). This assumption of the existence of neural attractor dynamics has formed the basis for multiple recent theoretical models of psychedelic’s effect on brain activity (Girn et al., 2023; Hipólito, Mago, Rosas, & Carhart-Harris, 2023; Ruffini, Lopez-Sola, Vohryzek, & Sanchez-Todo, 2024). In our context, these attractors corresponding to stable optima in the neural manifold of the PFC and represent task-relevant neural responses in a given environmental context (Beer, Lombardo, & Bhanji, 2010; Sarafyazd & Jazayeri, 2019). We interpret these attractors along the neural manifold as the priors or encoded beliefs which psychedelic-assisted psychotherapy holds potential in altering (Letheby, 2021; Zeifman et al., 2022).

Following these premises, we propose an optimization algorithm within the PP framework, which utilizes an energy-based model (EBM) of neural responses (Ackley, Hinton, & Sejnowski, 1985; Hinton, 1999; LeCun, Chopra, Hadsell, Ranzato, & Huang, 2006), whereby the neural response model attempts to predict incoming information from lower or upstream networks. In this perspective, the formation of attractors represents the persistence of stable responses, directions of change in neural responses can be modeled as gradients in the model’s energy landscape, driving the system towards stable states or trajectories corresponding to specific cognitive processes or behaviors. These attractors are equivalent to internal representations of beliefs from a cognitive perspective. Given the neural response variability observed in experimental studies, it has been proposed that neural responses might represent Monte Carlo (MC) (Metropolis, 1987) samples from a probabilistic response model, conditioned on stimuli (Hoyer & Hyvärinen, 2002; Orbán, Berkes, Fiser, & Lengyel, 2016). In this view, the neural response variability captures the uncertainty about the world inherent in the stimulus and context.

### 1.5 Operationalizing serotonergic neuromodulation

Starting from the theoretical models of Carhart-Harris and Nutt and Shine et al., we can understand the neuromodulatory role of serotonin in the cortex to be in service of improving computational efficiency. Within the EBM framework, computational efficiency corresponds to the ability to quickly sample accurate and predictive neural responses. We therefore concretely operationalize 5-HT2a and 5-HT1a agonism’s effect on the inference procedure through a transient modulation of the underlying energy function landscape. Although the relative density of these receptor populations varies throughout the cortex (Beliveau et al., 2017), we focus here on an abstract model of PFC dynamics, a region in which the expression of these two populations are relatively balanced (Amargós-Bosch et al., 2004). As such, we weight their modulatory contribution to the model equally.

A concrete mechanism for 5-HT2a agonism should meet a number of criteria. First, it must be excitatory, as postsynaptic 5-HT2a agonism in pyramidal cells is excitatory. Secondly, it should be consistent with the theoretical constructs of “active coping” and “flux” (Carhart-Harris & Nutt, 2017; Shine et al., 2022). Thirdly, it should induce an increase in the diversity of neural responses, as seen in neuroimaging of serotonergic psychedelic effects (Carhart-Harris et al., 2014). We therefore model 5-HT2a agonism as an ongoing stochastic perturbation of the energy landscape used to infer the neuronal response. This is an excitatory effect which actively encourages competition and encourages sampling novel neural responses which may have previously been difficult or impossible to reach in the original energy landscape. This increased sampling diversity comes at the cost of inducing a potential mismatch between the global optima in the modulated and unmodulated energy functions, an effect we will analyze below.

A concrete mechanism for 5-HT1a agonism should also meet certain criteria. Given the inhibitory postsynaptic effect of 5-HT1a agonism of pyramidal cells, it should be inhibitory. It should also be consistent with the constructs of “passive coping” and “automaticity” (Carhart-Harris & Nutt, 2017; Shine et al., 2022). We therefore model this effect as a transient global smoothing of the energy function landscape used to infer neural responses. This is an inhibitory effect, as it reduces the precision of the neural responses which are sampled. Unlike the 5-HT2a effect, it is also passive or automatic in that it does not change the relative energy assigned to different attractor points in the energy function landscape. It does however reduce the number of local minima in the landscape, moving the system from a highly competitive regime by increasing stability.

We recognize that while there is a theoretical basis for these modeling choices, the proposed computational mechanisms for 5-HT2a and 5-HT1a effects are novel and have yet to be empirically validated. Given the complexity of neuromodulation in the brain, there are other classes of receptor populations expressed throughout the brain which may serve similar computational roles in their impact on neural response sampling. As such, within the model and results presented below we refer to these mechanisms more generically as “excite” and “inhibit”, respectively. Based on the results of the model, in the discussion section we further elaborate on our hypothesized link between these mechanisms and serotonergic function, and its potential implications for clinical outcomes in the context of psychedelic-assisted therapy.

## 2 Methods

The model we propose relies on EBMs to model neural responses. We first present the relevant preliminaries concerning this class of model, followed by a description of our algorithm for sampling from EMBs with serotonin modulation; We then introduce the metrics of interest considered in our simulation results presented in Section 3. For additional details on the theoretical model and relevant connections to similar models, see Appendix B. For additional details on the specific parameters used in the simulations, see Appendix C.

### 2.1 Energy-based model preliminaries

From a predictive processing perspective, the brain is modeling the distribution of possible interoceptive and exteroceptive sensory signals which can be encountered in a given environmental context. Concretely, at any moment in time an organism is receiving some target sensory stimuli **x** which is distributed according to some probability distribution *p*(**x**). The goal of the model is to produce a neural response **z** which is capable of predicting **x** downstream.

Broadly, the distribution of stimuli is a function of some environmental context (Mante et al., 2013; C. Chen et al., 2024), which may persist for minutes, hours, days, or weeks (Muysers et al., 2024). This context may correspond to a particular configuration of social relationships, the physical environment, or goals which govern the individual’s mental and physical behavior. In psychedelic-assisted therapy, an environmental context would correspond to the “set and setting” for a particular drug-induced experience. In order to simplify the learning procedure, we consider a fixed environmental context, and correspondingly a stationary distribution over stimuli.

Pathologies in optimization correspond to failures to find stable and predictive configurations of neural activity, and likewise a failure to encode adaptive beliefs or internal representations. If these pathologies are severe and prolonged enough, they may lead to downstream failures in behavioral fitness and the development of psychopathology associated with either over-or under-canalization (Carhart-Harris et al., 2022; Juliani et al., 2024; Ruffini et al., 2024). These optimization issues can include the inability to find global optima (J. Chen & Luss, 2018; Ajalloeian & Stich, 2020), ill-conditioning of the energy landscape manifesting in high sensitivity in response to small changes in the input, deceleration in convergence to a stable attractor, overfitting and plasticity loss (Ahmed, Roux, Norouzi, & Schuurmans, 2019; Mei et al., 2020; Bolland, Louppe, & Ernst, 2023; Lyle et al., 2023).

In order to learn a neural response model, we take inspiration from control theory and machine learning (LeCun et al., 2006; Lotter, Kreiman, & Cox, 2016; Ranzato, Poultney, Chopra, & Cun, 2006; Ngiam, Chen, Koh, & Ng, 2011; Bond-Taylor, Leach, Long, & Willcocks, 2021), and employ EBMs, which perform inference and learning implicitly through an energy function, following prior approaches from the literature (Hoyer & Hyvärinen, 2002; Hennequin, Aitchison, & Lengyel, 2014; Aitchison & Lengyel, 2016; Orbán et al., 2016; Echeveste, Aitchison, Hennequin, & Lengyel, 2020; Qi & Gong, 2022; Dong & Wu, 2023). We define an EBM *p*(**z**) using a Boltzmann distribution as

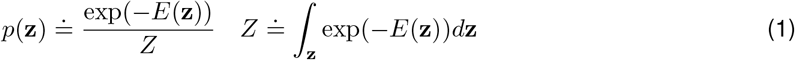

where *E*(**z**) is an energy function that maps input **z** ∈ *Ƶ* to a scalar, and *Z* is the normalizing constant with respect to **z** (also known as the partition function). Ideally, an energy function should assign low energy values to the samples drawn from data distribution, and high values otherwise.

In computational models of learning in the brain using EBMs, inference is carried out through neural sampling (Ackley et al., 1985; Hoyer & Hyvärinen, 2002), while learning happens via long-term synaptic plasticity (Bliss & Lomo, 1973; Miles, Poncer, Fricker, & Leinekugel, 2005). The neural sampling perspective of probabilistic inference in the cortex posits that the brain infers a distribution over neural responses consistent with Bayesian inference (Knill & Richards, 1996; Ernst & Banks, 2002; Körding & Wolpert, 2004; Knill & Pouget, 2004; Tenenbaum, Griffiths, & Kemp, 2006; Doya et al., 2006). To model the variability in neural responses, it has been proposed (Hoyer & Hyvärinen, 2002) to interpret the neural responses as Monte Carlo sampling (Metropolis, 1987) on the distribution *p*(**z**).

Prior works (Hoyer & Hyvärinen, 2002; Welling & Teh, 2011; Buesing, Bill, Nessler, & Maass, 2011; T. Chen, Fox, & Guestrin, 2014; Hennequin et al., 2014; Aitchison & Lengyel, 2016; Orbán et al., 2016; Echeveste et al., 2020; Qi & Gong, 2022) have developed and applied a number of approaches to sample from a distribution efficiently by defining the inference dynamics as performing walks in the latent space of neural responses, e.g., Gibbs sampling (Ackley et al., 1985), Langevin sampling (Neal, 2012; Welling & Teh, 2011; Genkin, Hughes, & Engel, 2021), Hamiltonian sampling (Neal, 2012; T. Chen et al., 2014). In this work, we are primarily interested in studying the effect of exogenous neuromodulation on the inference procedure of neural responses in EBMs, for which we rely on Langevin sampling.

For learning the energy function *E*(**z**) underlying the neural response model *p*(**z**), we follow prior research emphasizing the role of gradients in neural computation (Richards et al., 2019), and use a gradient-based learning algorithm, which corresponds to learning an energy function *E*(**z**) such that the distribution over neural responses under the model *p*(**z**) matches as accurately as possible a target distribution *p*^∗^(**z**).

To model neuromodulation, we introduce an auxiliary energy model, which is able to reflect signal enhancement caused by potential positive feedback loops that can occur under selective inhibition or excitation in latent circuit models (Aitchison & Lengyel, 2016). The resulting dynamical system is better able to explore the density and escape from local minima. The auxiliary energy, described in detail momentarily, is modeled as adaptively accumulating the neuromodulatory effects stemming from excitation-inhibition and homeostatic plasticity.

In order to keep our analysis clear and tractable, we make a set of simplifications to the PP framework. (i) First, we omit directly representing the stimuli **x**, assuming a supervised target distribution *p*^∗^(**z**) that the model is trying to learn *p*^∗^(**z**) ∝ exp(−*E*^∗^(**z**)). This is to provide an estimate of the fitness of a given neural response while avoiding the need to explicitly model the generation of stimuli. As such, *p*^∗^(**z**) corresponds to a distribution over neural responses which is able to optimally predict the stimuli distribution downstream. (ii) Second, *E*(**z**) and *E*^∗^(**z**) are both modeled as a tabular representation, with *N* × *N* set of points, each with an associated value that can be changed during learning. Thus, the synaptic connectivity is updated directly through this tabular representation. A consequence of the tabular representation of *E*(**z**) is that all gradients in the simulator are approximated using finite differences.

### 2.2 Concrete algorithm for neuromodulated EBMs

The learned and target energy functions *E*(**z**) and *E*^∗^(**z**) are sampled from a distribution of possible energy function landscapes *E* ∼ *ε* (*N, σ*). Specifically, *ε* (*N, σ*) = *M* ∗ *G*(*σ*), where *M* ∈ ℝ^*N ×N*^ with *M*_*ij*_ ∼ *𝒩* (0, 1), ∀*i, j* ∈ {1, …, *N* }, *N* is the size of the tabular representation, *G*(*σ*) is a Gaussian function which is convolved with *M*, and *σ* is the size of the Gaussian function kernel. The values of these representations are then normalized to be within the range [0, 1] at the start of the optimization process.

Finally, we employ a simplified learning objective for *E*(**z**) ∝ − log *p*(**z**) to describe changes in synaptic connectivity, namely via min_*E*_ ℒ (*E*), with 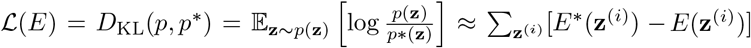, capturing the difference in energy function values at sampled responses. These neural responses **z** are sampled from an auxiliary energy function 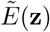, a surrogate corresponding to the energy function under neuromodulation.

#### Neuromodulatory signals

We consider two unique neuromodulatory signals and one additional constraint:

- *α*^excite^: modulates the excitatory drug effect
- *α*^inhibit^: modulates the inhibitory drug effect
- *α*^homeo^: modulates the strength of homeostatic constraints

#### Sampling neuromodulation level

The ranges for *α*^excite^ and *α*^inhibit^ can either be set constant for the duration of the optimization process, or be determined dynamically. Given that we are interested in exogenous neuromodulation effects from a drug, we use a function `drug` (*t*) to determine the neuromodulation levels as a function of time. This function uses the probability density function of a beta distribution to provide a response curve used to determine the drug strength over time. This function is used to provide a simple and generic representation of prototypical change in blood plasma concentration levels of a drug in which there is a rapid increase in concentration followed by a slower decrease in concentration as the drug is fully metabolized.

##### Algorithm 1 Neuromodulated optimization algorithm

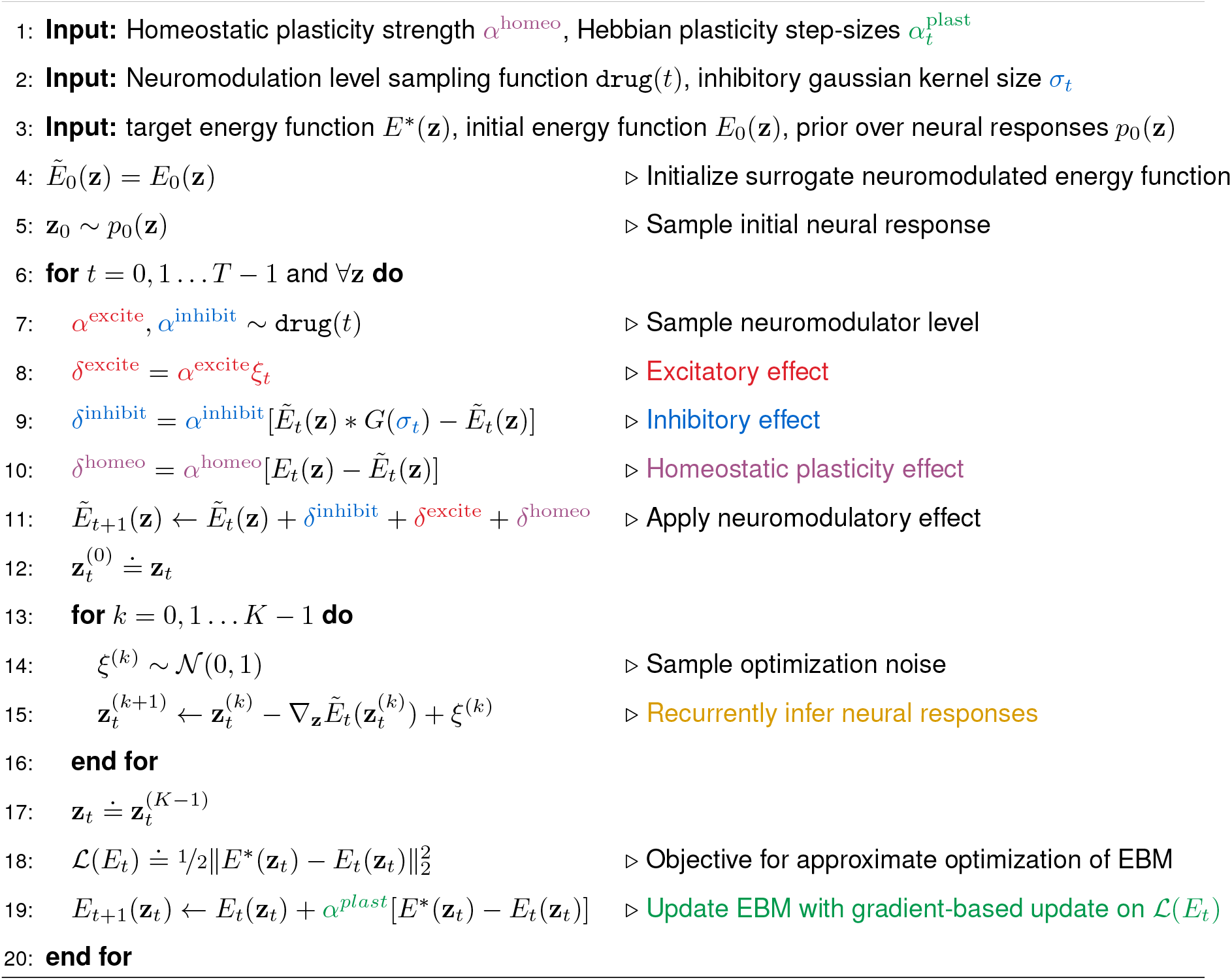

#### Excitatory drug effect

At each iteration *t*, a stochastic perturbation is applied to the auxiliary energy function 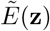

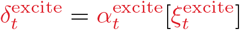

where 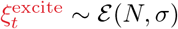 is structured noise sampled from the energy function distribution and then scaled to be within the range [−0.5, 0.5], and 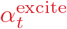 the relative excitatory strength of neuromodulation.

#### Inhibitory drug effect

We simulate the inhibitory effect by applying a Gaussian filter kernel centered around 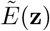 and apply this effect to the auxiliary energy function 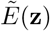

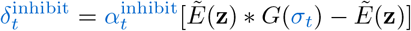

We use a local Gaussian filter kernel *G*(*σ*) which is convolved with 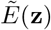, where *σ*_*t*_ denotes the size of the Gaussian kernel and 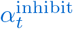 the relative inhibition strength of neuromodulation.

#### Homeostatic plasticity effect

Homeostatic plasticity is a mechanism for stabilizing neural activity levels across a network, ensuring that neurons maintain an appropriate level of excitability. This regulatory mechanism restores stability and mitigates imbalances in the activity levels of neurons by regularizing the neuromodulated energy landscape towards the baseline

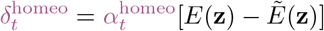

with 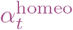 the relative strength of the update toward the previous value of the unmodulated energy function *E*(**z**).

#### Applying neuromodulatory effects

The three neuromodulatory effects are then additively combined into a single term *δ*_*t*_ and accumulated into 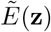 before the sampling process takes place each iteration. The optimization procedure including the aforementioned neuromodulation mechanisms is described in Algorithm 1, and visualized in Figure 3.

**Figure 3.**
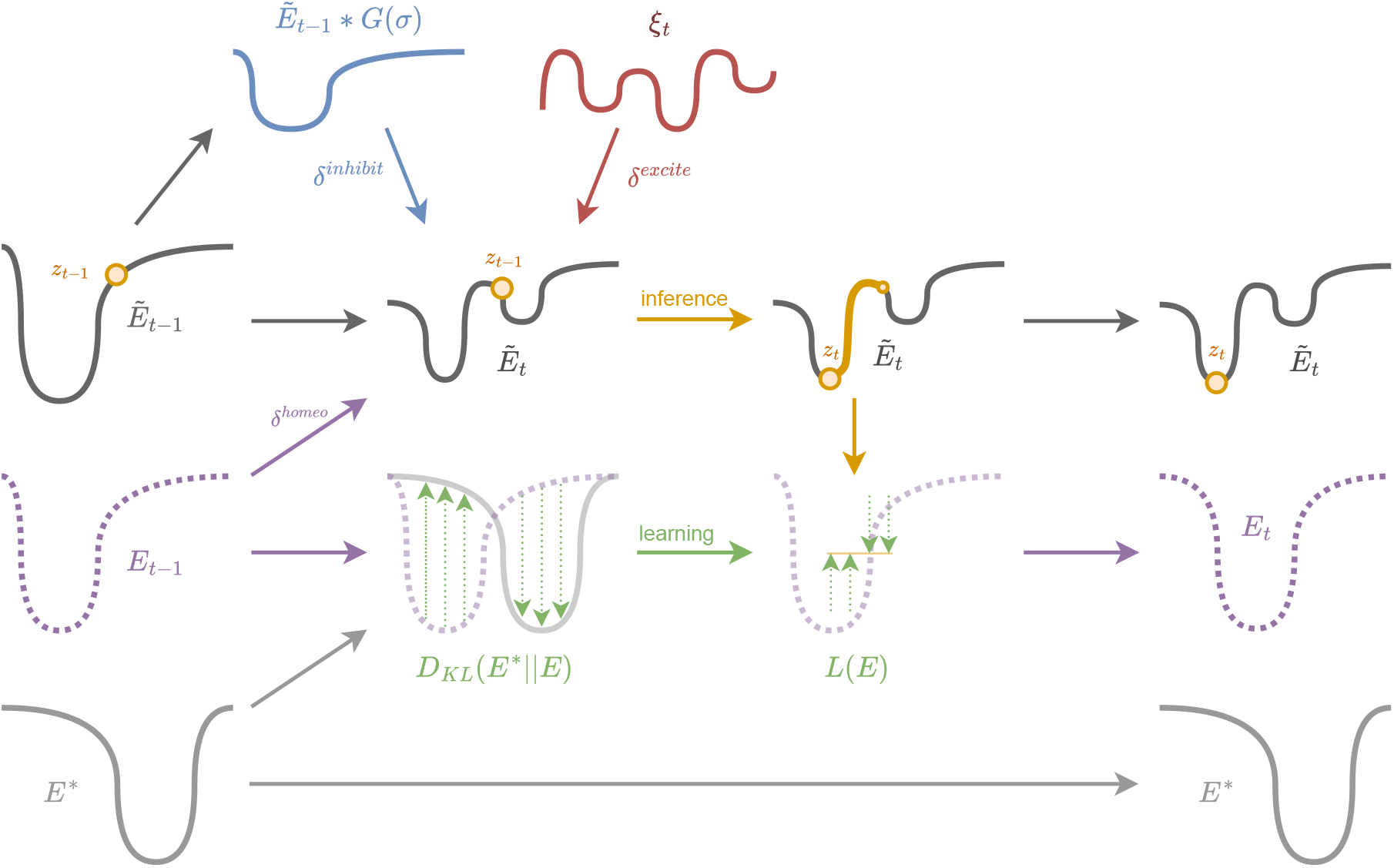
Simplified diagram of the inference and learning procedure used to update the internal representation **z**, the energy function *E*_*θ*_(**z**), and the neuromodulated energy function 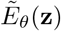 at each time-step, with time moving from left to right. A one-dimensional energy function is used here for illustrative purposes. Top row displays the generation and application of excite *ξ*_*t*_ and inhibit 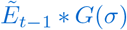 modulation, along with the homeostatic effect from *E*_*θ*_(**z**) to the neuromodulated energy function 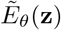. Second row displays the neuromodulated energy function 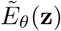 undergoing modulation, followed by the inference procedure used to sample *z*_*t*_ taking place. Third row displays the non-modulated energy function *E*_*θ*_(**z**) being updated based on error signals *L*(*E*) generated from the inference procedure against the target function *E*^∗^(**z**), represented in the bottom row as unchanging. See Algorithm 1 for a more detailed account of the inference and learning process.

### 2.3 Metrics of interest

We consider six primary metrics of interest which we collect and analyze during each simulation experiment. Figure 4 (1st row) contains a simple graphical representation of each metric under consideration, with the final two being represented by the rightmost graphic.

**Figure 4.**
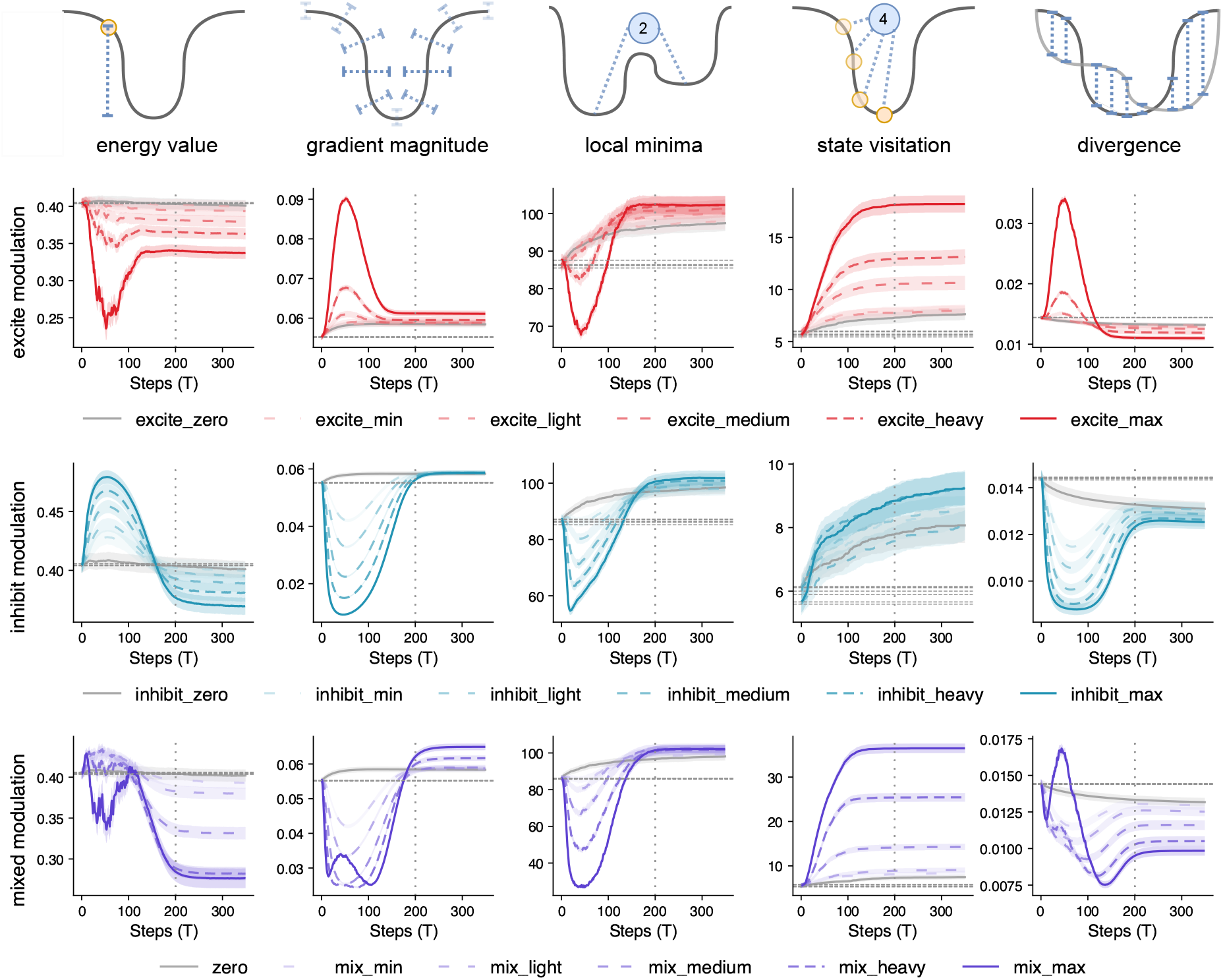
**(First row)** Graphical representations of the five primary metrics considered in the simulations. The representation in the header of each column is simplified to use a one-dimensional energy function for visualization purposes whereas the energy function used in the simulation is two-dimensional. The representation corresponds to the neuromodulated energy function 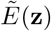. Orange circles correspond to samples from the EBM 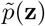. For each metric we convey in blue an intuitive depiction of its measurement. The x-axis shows the number of steps *T* in Alg.1. **(Second row)** Shows the average metric values for excitatory modulation experiments. **(Third row)** Shows the average metric values for inhibitory modulation experiments. **(Fourth row)** Shows the average metric values for balanced mixed modulation experiments. In all time series plots the solid grey line corresponds to the baseline without neuromodulation. Horizontal dashed line represents value of metric at start of drug intervention. Vertical dotted line represents point at which drug effect ends. Semi-transparent bands around lines correspond to standard error.

#### Energy function value

The energy function value corresponds to the 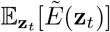 for a set of sampled **z**_*t*_ values at a given iteration *t* of the simulation. This metric provides insight into how a given intervention on the neuromodulatory system changes the energy level of sampled **z**_*t*_ values relative to a non-modulated baseline over time. The EBM represents the probability of different responses implicitly, by the frequency with which it visits their representations via its dynamics. Higher energy of some neural response **z** means neural activity will likely visit **z** with less frequency, whereas lower energy corresponds to a larger proportion of the time neural activity representing **z**.

#### Gradient magnitude

We consider the average gradient magnitude of the energy function at a given time-step. This is calculated by uniformly evaluating 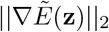, with ∥ *·* ∥_2_ the *L*_2_ norm, for every possible value of **z** in our tabular representation. Functions with bounded gradient magnitudes are smooth, ensuring that changes in input variables result in proportionally limited changes in the function’s output. This is crucial for maintaining stability and convergence of optimization algorithms towards optimal solutions. The average gradient magnitude value also corresponds to the precision assigned to a neural response in the competitive motif. The magnitude of the gradient is correlated with the stationarity of the environmental context. Larger magnitudes indicate sharp changes in the energy function and are desirable only to the extent that the neural responses **z** are optima and both able to reduce prediction errors in the current and future environmental contexts. This is more likely to be the case the more stationary the environmental context is over time. In non-stationary environments, a smaller average gradient magnitude over time will correlate with smoothness, and may be desirable as it will produce a more diverse sampling of **z**.

#### Local minima count

We consider the number of local minima present in the energy function at a given time-step. This is calculated by uniformly evaluating our tabular representation of 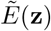 and determining the local gradient at each point. The local minima count reflects the complexity of the energy landscape (the more local minima, the more complex the landscape) and corresponds to possible stable attractor states in which the stochastic sampling process may enter. Within the context of our theoretical model, the number of local minima is predictive of how successful both the learning and inference processes will be. Fewer local minima increase the likelihood that the stochastic sampling process will infer a neural response closer to global minima of the energy function.

#### State visitation count

We calculate state visitation count by tracking the number of unique values which **z** takes over the course of the optimization process in a given experiment. Within the context of our theoretical model, this metric illustrates the ability to cover a broad range of plausible interpretations of its input stimuli by exploring the entire posterior space, where better coverage (i.e. more visited states) leads to more accurate estimation of the energy function. As such, this metric can be seen as a proxy for diversity over neural states. The number of states **z** is correlated with diversity of neural activity patterns which might be recorded using neuroimaging techniques. Given the inherent low-dimensional nature of our **z** values, we simply count unique states rather than compute typical complexity measures such as Lempel–Ziv, which assumes the compressibility of states.

#### Energy function divergence

We also track the surrogate objective 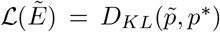, where 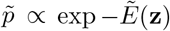, defined as the KL-divergence between the posterior probability distribution induced by the auxiliary variable—a surrogate energy function 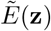 resulting from the acute drug-infused neuromodulation, and the target distribution induced by *E*^∗^. This value is a proxy for the magnitude of the prediction error that can be expected by sampling from 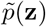 at a given iteration, and thus smaller values correspond to reduced expected error. Generally, an optimal probabilistic model over neural responses (i.e. 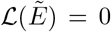) would imply that an organism is able to infer stimuli correctly and reduce uncertainty relative to its environment.

#### Divergence trend monotonicity

Finally, we consider a measure that quantifies the evolution of the learning objective or regret—the KL-divergence loss, during optimization. In an ideal scenario, the convergence curve of the optimization process would show a consistent decrease over iterations. However, in practice, there exist fluctuations or plateaus due to system noise and the influence of the neuromodulatory signal. Consequently, we resort to quantitatively capturing the overall compressed learning trend via the cumulative increase in regret over the course of an experiment. This corresponds to the cumulative sum of positive consecutive sub-optimality gaps, measured under the KL-divergence _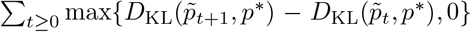_ evaluated between each pair of consecutive iterations. As a result of the learning process, the learning loss captured by the KL-divergence should decrease over time, so the cumulative regret should be negative 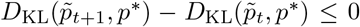. This metric quantifies fluctuations and transitory changes, thereby measuring the disruption in the learning process caused by neuromodulation. Within the context of our theoretical model, this corresponds to the extent to which uncertainty or misaligned neural responses are introduced by the pharmacological intervention.

## 3 Results

In order to evaluate our proposed energy-based model with neuromodulation, we run a series of idealized simulations of the algorithm presented above. We first examine the individual impacts of excitatory and inhibitory neuromodulation on cortical optimization, finding that they respectively induce transient overfitting and underfitting in the optimization process of the posterior probability over neural responses. We next consider the effect of mixed neuromodulation, and find that this induces complex synergistic effects which retain desirable properties of each individual form of modulation. Finally, we explore the full space of biased neuromodulation and report the relative level of excitation and inhibition which optimally trades off between our proxy metrics, finding it to be a biased inhibitory neuromodulation.

### 3.1 Excitatory neuromodulation induces transient overfitting

We first consider the effects of pure excitatory modulation on the final KL-divergence values across a range of dose levels (see Figure 4, 2nd row). We find that only two levels of 5-HT2a agonism decrease the KL-divergence relative to the baseline: heavy [two-tailed t-test: *t*(99) = 3.1, *p <* 0.001], and max [*t*(99) = 4.83, *p <* 0.001] (column 5).

When we examine the energy function values over the course of the experiment, we observe that across all dose ranges there is a transient acute reduction of the energy value below the baseline. Furthermore, in cases of `heavy` and `max` dose levels, this acute reduction falls below the final values in the post-acute stage of the simulation (column 1). We also find that the gradient magnitude increases over baseline during the acute modulatory effect, and that this effect is likewise correlated with dose strength, with `heavy` [*t*(99) = −3.249, *p <* 0.001] and `max` [*t*(99) = −11.391, *p <* 0.001] doses inducing significantly larger gradient magnitudes during the acute effect (column 2).

The transient increase in gradient magnitude paired with the sampled energy value becoming lower than would be possible in the absence of neuromodulation can be interpreted as the induction of transient overfitting in neural response inference during the acute phase of excitatory modulation when a dose strength of `heavy` or `max` is applied. Significantly, this overfitting is not to the target distribution *p*^∗^(**z**), but rather to the distribution induced by the current neuromodulated energy function 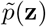. This transient overfitting may serve as a basis for the induction of transient (and sometimes false) insight experiences and supernatural percepts during the acute effects of psychedelic drugs (Griffiths, Hurwitz, Davis, Johnson, & Jesse, 2019; McGovern et al., 2023).

Turning to the remaining metrics, we observe that local minima count decreases relative to the baseline during the acute phase of neuromodulation for the `heavy` and `max` dose levels [both *p <* 0.001] (column 3). State visitation count in contrast increases according to the dose strength, with `medium`, `heavy`, and `max` doses producing significantly higher state counts than baseline [all *p <* 0.005] (column 4). Finally, we find that as the dose strength increases the KL-divergence metric increases as well, with significant differences found for all levels [all *p <* 0.001] (column 5).

As a result of excitatory modulation, both the number of attractor states and the location and energy value of these states change significantly during the acute modulatory phase and this effect is correlated with modulation strength. These results suggest the number of attractor states decreases and the optimization procedure exhibits oscillatory or diverging dynamics during the acute modulation phase, an effect correlated with modulation strength. This finding is consistent with observed increases in the entropy of cortical dynamics as measured by fMRI in individuals experiencing the acute effects of serotonergic psychedelics (Carhart-Harris et al., 2014).

### 3.2 Inhibitory neuromodulation induces transient underfitting

We next consider the effects of pure inhibitory neuromodulation on the metrics of interest across the same range of doses (see Figure 4, 3rd row). We first find that inhibitory modulation is unable to produce a lasting decrease in the KL-divergence between the learning and target posterior probability distributions, even at the `max` dose level (column 5). On the other hand, during the acute phase of the simulation we find that inhibitory modulation transiently, but significantly decreases the KL-divergence values for all levels of modulation [all *p <* 0.001] (column 5).

Similar to what we observe with excitatory modulation, we find that inhibitory modulation is able to induce decreases in the post-acute energy function values. We find however that only the max dose level is able to produce a significant decrease compared to the baseline [*t*(99) = 3.498, *p* = 0.001] (column 1). Unlike excitatory modulation, inhibitory modulation induces an opposite effect during the acute phase by increasing the energy value above baseline in a manner correlated with modulation strength, with `medium`, `heavy`, and `max` doses producing significant acute increases [all *p <* 0.001]. Examining the change in gradient magnitude, we find an inverse effect to that of excitatory modulation, with inhibitory modulation inducing significant transient decreases in gradient magnitude as a function of modulation strength for all dose levels [all *p <* 0.001] (column 2).

Increases in energy values along with decreases in gradient magnitude together can be interpreted as the induction of transient underfitting in neural response inference procedure during the acute effects of inhibitory modulation. As with the excitatory effect, this underfitting is not to the target distribution *p*^∗^(**z**), but rather to the distribution induced by the current neuromodulated energy function 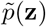. This transient underfitting is equivalent to belief relaxation, where the precision of encoded beliefs is significantly decreased. These transient effects are also correlated with the acute decrease in the KL-divergence objective metric.

Examining the remaining metrics, we find that inhibitory modulation produces a significant acute decrease in the number of local minima within the energy function which are greater than those of excitatory modulation (column 3), and that this effect is correlated with dose strength [all dose levels above min *p <* 0.001]. We find that the number of visited states is also significantly increased by inhibitory modulation (column 4), though at a level which is considerably less than that of excitatory modulation, with only `max` [*t*(99) = −2.104, *p* = 0.037] dose levels produce significantly greater state counts than baseline. These results suggest that inhibitory modulation alone is unlikely to produce the increases in neural response diversity comparable to those which result from administration of serotonergic psychedelics (Carhart-Harris et al., 2014).

### 3.3 Complex effects of balanced mixed neuromodulation

We next report the effect of applying evenly balanced excitatory and inhibitory modulation on the metrics of interest across a range of doses (see Figure 4, 4th row). We find that these neuromodulation patterns are able to produce final KL-divergence values which are significantly lower than the considered baseline for the `medium`, `heavy`, and `max` dose levels [all *p <* 0.001] (column 5). We also observe that for all levels except for `max`, there is a transient decrease in KL-divergence values during the acute phase of modulation. Balanced modulation also produces significant decreases in post-acute energy function values at the `medium`, `heavy`, and `max` dose levels [all *p <* 0.001] (column 1). Furthermore, these values are lower than either pure excitatory or inhibitory modulation alone at the equivalent dose level. The acute energy function values for balanced agonists are also neither above or below pre-or post-acute levels. We also observe that mixed modulation produces significant transient decreases in gradient magnitude at all dose levels [all *p <* 0.001], but that these are less extreme than those of equivalent pure inhibitory modulation.

Taken together, these results suggest that balanced modulation is able to reduce post-acute energy function values without the introduction of either predominant underfitting or overfitting of the neural response function during the acute phase. Given the transient but unstable decrease in gradient magnitude, we can still interpret these interventions as inducing a form a belief relaxation, although one which is not as consistent as that of pure inhibitory modulation. It also differs from pure inhibitory modulation in that the states being sampled have a higher likelihood than they would under the baseline (as a result of the lower energy value), but a lower precision (as a result of the lower average gradient magnitude).

Turning to the remaining metrics, we find that balanced modulation induces a decrease in the number of local minima during the acute drug phase which is greater than that seen for either excitatory or inhibitory modulation alone [all except `min` dose level *p <* 0.001] (column 3). Correspondingly, state visitation count is also increased in balanced modulation over excitatory or inhibitory modulation, with all dose levels above `min` producing significantly more visited states [*p <* 0.01] (column 4). Relative to monotonic decrease of the optimization objective over the course of the simulation observed in the baseline, here we observe that there are significant increases in KL-divergence [all *p <* 0.001] (column 5). Notably however we see more monotonicity in the KL-divergence trend compared to what is induced by excitatory modulation.

### 3.4 Characterizing the space of biased neuromodulation

Above we found a synergistic effect of balanced modulation on post-acute values of the KL-divergence as well as the state visitation and acute local minima counts. Balanced modulation also produced a moderating effect on the non-monotonicity of the KL-divergence trend. Given these improvements as compared to pure excitatory or inhibitory modulation, we hypothesized that a neuromodulation profile which is biased rather than balanced may prove even better at minimizing KL-divergence while retaining a largely monotonic divergence trend. In order to determine this, we simulated all possible permutations of excitatory and inhibitory modulation strengths. These results are presented in Figure 5.

**Figure 5.**
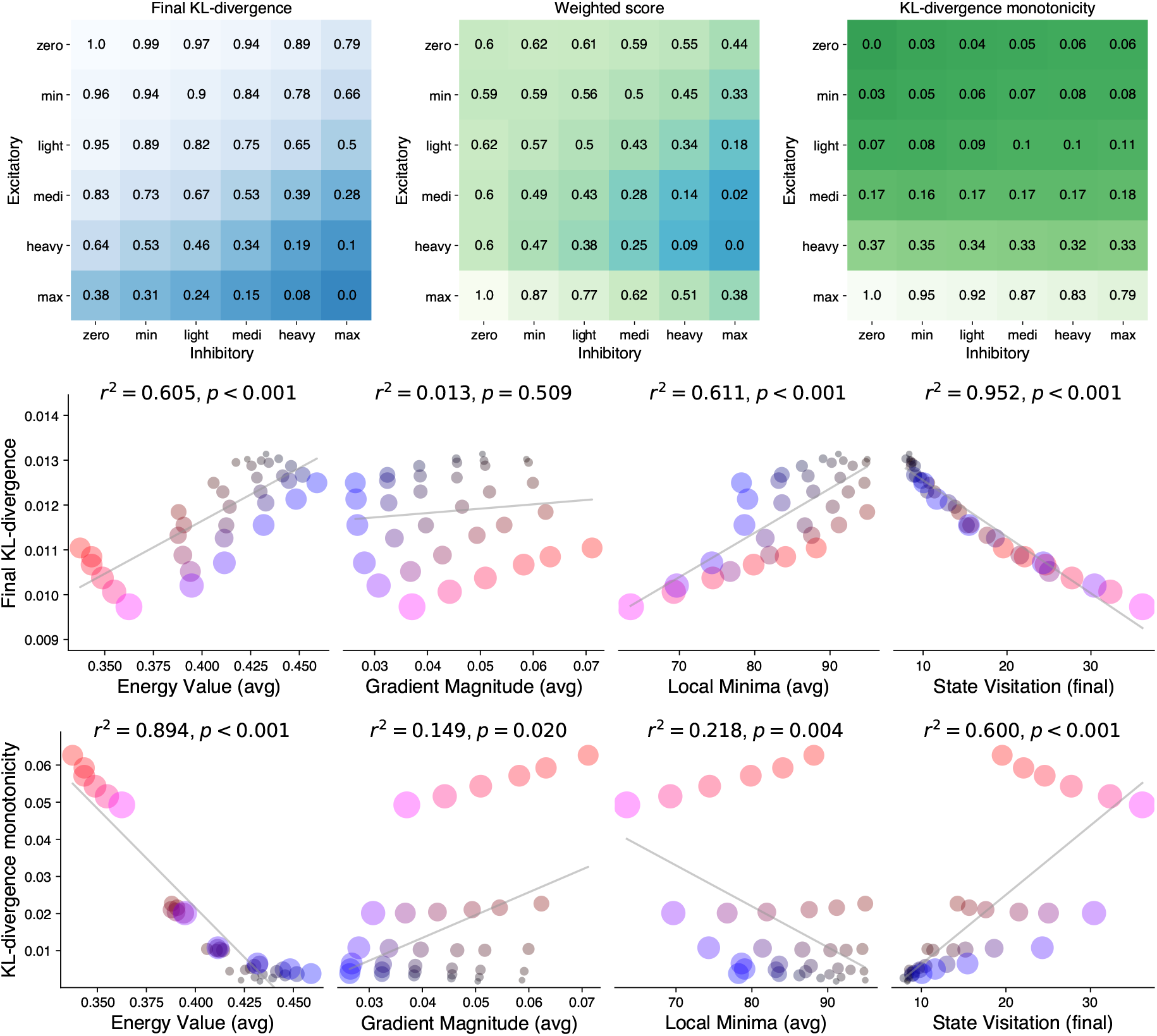
**(Top-left)** Matrix reporting normalized final energy values for set of possible excitatory and inhibitory modulation combinations. Lower values correspond to lower energy. **(Top-right)** Matrix reporting normalized final non-monotonicity values for set of possible excitatory and inhibitory modulation combinations. Lower values correspond to higher monotonicity. **(Top-middle)** Matrix reporting normalized weighted energy values for set of possible excitatory and inhibitory modulation combinations. Lower values correspond to lower weighted score. **(Middle)** Correlations between metrics and final KL-divergence values. **(Bottom)** Correlations between metrics and KL-divergence monotonicity. Color of data points correspond to relative excitatory (red) and inhibitory (blue) modulation. Size of data point corresponds to average modulation strength level.

We find that a `max` dose of a balanced excitatory and inhibitory modulation produces the greatest post-acute reduction in KL-divergence. Likewise, the general trend is that increases across both modulator ranges produces improvements in this metric, with a greater portion of the improvement coming from increases in excitatory rather than inhibitory modulation. In contrast, we find that the KL-divergence trend is most monotonic at low levels of both excitatory and inhibitory modulation, but is most strongly affected by increases in excitatory modulation.

In order to trade-off between these two metrics, we normalized values of both final KL-divergence and divergence trend non-monotonicity and equally weighted them to produce a new score metric which balances the two desirable outcomes. Here we find that a `max` level of inhibitory modulation paired with a `medium` or `heavy` level of excitatory modulation provides the optimal trade-off between the two metrics of interest. In particular, this level of biased modulation produces post-acute KL-divergence values which are comparable to that of pure max excitatory modulation, but does so with a trend which is significantly more monotonic than that produced by max excitatory modulation.

### 3.5 Relationship between properties of neuromodulation and outcomes

We finally consider the relationship between the metrics describing the acute modulation effects and our two proxy outcome metrics. We find that while average gradient magnitude is uncorrelated with the final KL-divergence, final energy value, average local minima count, and final state visitation count are all significantly correlated with final KL-divergence value [linear regression: all *p <* 0.001] (see Figure 5). Among these, state visitation count is able to account for nearly all of the variance in final optimization objective, with an *r*^2^ = 0.952.

Average energy value, average gradient magnitude, and final state visitation count are also linearly predictive of monotonicity in the KL-divergence trend [all *p <* 0.01]. Of these, we find that average energy value accounts for the greatest variance in the proxy metric, with an *r*^2^ = 0.894. Although this relationship is not causal, it does imply that overfitting in the energy function during the acute drug experience is associated with greater divergence between the current and target energy distributions.

## 4 Discussion

Operating within the preexisting theoretical frameworks of predictive processing and energy-based models, we proposed concrete mechanisms whereby 5-HT2a and 5-HT1a agonism drugs such as psilocybin, LSD, and DMT may exert their effects, and simulated these effects across a range of dose strength and relative agonism biases. Our model and results have a number of potential implications for debates around belief representation, clinical application of psychedelic-assisted therapy, as well as next-generation psychedelic drug design. Below we attempt to address these considerations.

### 4.1 Mechanisms of psychedelic-induced belief alteration

Our hypothesis is that the excitatory and inhibitory effects modeled above are useful proxies for the effects of 5-HT2a and 5-HT1a agonists on cortical dynamics (see Section 1.5). Here we take up the implications of this connection in light of the results from our model. We first consider the debate around whether psychedelics are belief-strengthening, belief-relaxing, or some third category (Carhart-Harris & Friston, 2019; Safron, 2020).

The results of our model indicate that 5-HT2a agonism can be characterized as acutely inducing a transient overfitting into the cortical dynamics, resulting in the generation of a sequence of ever-shifting high-precision beliefs which may or may not be veridical. Strictly speaking, we can interpret this transient overfitting as a form of acute belief-strengthening. Rather than a fixed set of beliefs being strengthened for the duration of the drug effect, the increase of belief precision is stochastic and transient, resulting in a higher state visitation count as attractors in the energy landscape are constantly disrupted during the acute drug effect. As such, this strengthening of beliefs does not refer to post-acute outcomes, and is therefore compatible with findings that post-acutely serotonergic psychedelics result in more relaxed beliefs (Zeifman et al., 2022).

These transient high-precision beliefs may correspond to insight experiences, a common characteristic of the phenomenology of serotonergic psychedelics (Laukkonen et al., 2023). These experiences are often characterized by a “noetic feeling,” or sense of acquired knowledge through an unknown or mystical source (Hewitt, 2011). While these experiences may contribute to therapeutic outcomes, by overfitting to the neuromodulated energy function, these insights may be “false” in that they are not reflective of an actual capacity to better predict neural responses in a non-modulated state (McGovern et al., 2023). Likewise, such false insights are often discarded or forgotten post-acutely due to their non-veridical nature or in-applicability to the current behavioral context the individual finds themselves within.

Our model predicts that 5-HT1a agonism also significantly contributes to the belief-altering properties of serotonergic psychedelics by inducing a belief-relaxation effect, which can be measured both by the decrease in gradient magnitude as well as energy function values during the acute phase of the experiment. Both of these metrics together serve as our proxies for belief precision, suggesting that our models predicts that 5-HT1a agonists will acutely produce belief relaxation. This can be characterized as acutely inducing transient underfitting into cortical dynamics, resulting in the generation of a more stable sequence of lower-precision beliefs than either a placebo baseline or 5-HT2a agonism.

Also consistent with belief relaxation is our model’s prediction that 5-HT1a agonists decrease the number of local minima present in the energy function surface, which corresponds to the number of stable beliefs a network can encode. This constellation of effects is consistent with the phenomenological reports of 5-MeO-DMT use, a highly biased 5-HT1a agonism which at heavy doses produces experiences in which users encounter a “white-out” state in which they remain conscious but become unable to represent objects, ideas, space, or even time (Reckweg et al., 2022; Dourron et al., 2023; Gómez-Emilsson, 2020).

Our model predicts that mixed agonists such as psilocybin, LSD, or DMT will produce a more complex neural signature. In this case we find both an increase in state visitation count as well as an overall decrease in gradient magnitude and number of local minima. Furthermore we find that underfitting is more apparent for most of the dose-response curve, but overfitting is induced at the high end of the curve, at which point the 5-HT2a effect predominates. This predominance of 5-HT2a over 5-HT1a at high levels of agonism is characteristic of endogenous serotonergic effects on prefrontal cortex dynamics (Puig & Gulledge, 2011), and part of the “flux” vs “automaticity” model (Shine et al., 2022).

This opponency effect between 5-HT2a and 5-HT1a agonism on belief representation has previously been identified in the context of a model contrasting DMT and 5-MeO-DMT (Gómez-Emilsson, 2020). It is also is consistent with the passive vs active coping model introduced by Carhart-Harris and Nutt. In all cases, 5-HT1a agonism can be understood to acutely relax beliefs whereas 5-HT2a agonism acutely and stochastically strengthens beliefs. The effect of the former is to produce a state which may provide short-term relief from stress, as is seen in the anxiolytic effects of 5-HT1a agonists. The latter in contrast provides the potential for long-term benefit. Taken together, these findings are both consistent with the ALBUS and REBUS models of psychedelic action. Whether psychedelics are viewed as strengthening or relaxing beliefs is then a question of the particular drug, timescale, and proxy metric of interest under consideration.

### 4.2 Potential implications for clinical research

Error minimization in the brain is ultimately in service of the long-term behavioral success of an organism (Kirchhoff, Parr, Palacios, Friston, & Kiverstein, 2018). Theoretical work has linked this success in minimizing prediction errors to positive mental health outcomes (Smith, Badcock, & Friston, 2021), and conversely the chronic failure of predictive error minimization to negatively valenced affect (Smith et al., 2022). Greater felt uncertainty for example is often characteristic of anxiety and mood disorders (Carleton et al., 2012; Smith, Kirlic, et al., 2021). The presence of systemic and persistent prediction errors in cortical representation reflect an agent’s inability to resolve the uncertainty in an environment. This may particularly be the case when the inability to resolve prediction errors takes place at higher levels of the cortical hierarchy, as this suggests a fundamental maladaptivity in the foundational beliefs of the individual.

Prediction errors in themselves are not pathological, as they may simply indicate the presence of novel information within the environment. Indeed, it is believed that exploratory behavior in animals is largely driven by the seeking of stimuli capable of producing prediction errors (Clark, 2018), and prediction errors can also correspond to better-than-expected behavioral outcomes (Schultz, 2017). Under healthy circumstances, these errors are quickly utilized to update beliefs and the errors are minimized. In the perspective discussed here, various forms of psychopathology may arise from a failure to appropriately update maladaptive beliefs, resulting in the presence of persistent and systemic prediction errors (Carhart-Harris et al., 2022).

Starting from these theoretical assumptions, the simulation results presented above contain multiple theoretical metrics which we believe can be interpreted as potentially relevant to clinical outcomes. We propose that the KL-divergence between target and current sampling distributions may serve as a proxy for the overall clinical efficacy of an intervention. The logic behind this connection is that the KL-divergence is a metric of the frequency and magnitude of systemic prediction errors. In our model, initial KL-divergences were large, due to the starting energy function and target functions being drawn as independent samples and thus de-correlated. Because the presence of systemic prediction errors is associated with psychopathology, reducing this metric should correspond to improvements in mental health outcomes.

We can make a speculative interpretation of our results using this framework. Our model predicts that biased 5-HT2a, but not 5-HT1a agonists are able to reliably reduce post-acute KL-divergence as compared to a placebo. We can interpret this as a prediction that acute administration of 5-HT2a agonists will be capable of producing long-term therapeutic outcomes, whereas acute administration of a 5-HT1a agonist will not. The model also predicts that mixed agonists produce greater post-acute reductions in KL-divergence than either biased agonist alone. Under this interpretation, mixed agonists such as LSD, DMT, and Psilocybin are predicted by our model to be more clinically effective in the long-term treatment of psychopathology than pure agonists such as DOI or 8-OH-DPAT.

The second metric of clinical relevance is the KL-divergence monotonicity over the course of the intervention. Because we interpret reductions in KL-divergence compared to baseline as a metric for the efficacy of the treatment, we can likewise interpret acute increases in this metric which may take place during the intervention as a proxy for the acute tolerability of the intervention. Our model predicts that biased 5-HT1a, but not 5-HT2a agonists are likely to be acutely tolerable. Likewise, the model predicts that mixed agonists are more tolerable than biased 5-HT2a agonists due to the modulating effect of 5-HT1a agonism. We find evidence of the modulating effect of 5-HT1a agonism in the literature, where again mixed agonists such as LSD and DMT have received the majority of the clinical attention for their favorable tolerability profile. This is also consistent with pre-existing findings that 5-HT1a agonists produce acute anxiolytic effects, while 5-HT2a agonists can be anxiogenic under certain circumstances. There already exists some evidence that increasing 5-HT1a receptor activation during the acute psychedelic experience decreases the intensity of the reported acute experience (Strassman, 1995; Pokorny, Preller, Kraehenmann, & Vollenweider, 2016), but this should be examined in greater depth by explicitly modulating the relative levels of 5-HT2a and 5-HT1a agonism across a range of doses.

Using these two hypothesized proxy metrics, we can also interpret the weighted scores which were generated for the sweep of mixed agonists with different biases. We found that overall score was highest for mixed agonists with max 5-HT1a and heavy/medium 5-HT2a. As such, our model predicts that serotonergic psychedelics with this 5-HT1a biased binding profile may provide an optimal trade-off between acute psychological tolerability and long-term clinical efficacy. This class of drugs would include DMT, 5-MeO-DMT, and other 5-MeO compounds such as 5-MeO-DALT. It is important to note that these interpretations of our model are speculative, and will need extensive testing and validation before they can be relied on to guide clinical decision making. Regardless, we believe they provide a potentially useful working model which can be used to generate testable (and falsifiable) hypotheses regarding clinical outcomes from serotonergic psychedelics.

/addedWe can also interpret the state visitation count as a proxy for the entropy of the cortical dynamics. Our model predicts that unlike 5-HT2a agonist psychedelics, 5-H1a agonists will not significantly increase acute cortical entropy, as measured by the number of unique cortical states visited as well as the transition dynamics between states (Carhart-Harris et al., 2014). Our model also predicts that balanced mixed agonists will produce levels of cortical entropy which are significantly greater than either 5-HT1a or 5-HT2a agonists alone, suggesting a complex synergistic effect between the two receptor systems. Furthermore, we find that increases in state visitation count are predictive of final KL-divergence, our proxy metric for therapeutic efficacy. This is consistent with Carhart-Harris et al., who also predict that increases in cortical entropy will be significantly correlated with greater post-treatment therapeutic efficacy.

### 4.3 Guiding a novel direction in drug design

Current efforts in psychedelic drug design are largely focused on the development of non-hallucinogenic 5-HT2a agonists (Cameron et al., 2021; Qu et al., 2023; Lewis et al., 2023). Although progress has been made in this area, there is evidence to suggest that the acute phenomenology of the psychedelic experience plays a non-trivial causal role in the long-term therapeutic efficacy of psychedelic-assisted therapy (Griffiths et al., 2011; Yaden & Griffiths, 2020). If this is indeed the case, then there is value in understanding the pharmacological mechanisms responsible for different aspects of the acute phenomenology and their relationship to clinical outcomes.

Early clinical evidence suggests that 5-MeO-DMT, a highly biased 5-HT1a agonist psychedelic, has significant potential as a treatment for major depressive disorder (Uthaug et al., 2019; Reckweg et al., 2023). Some concern exists however regarding the safety profile of 5-MeO-DMT both in the acute and post-acute phases of the drug effect. One particular concern is the experience of “reactivations” in which the drug phenomenology of the effect spontaneously returns days or weeks after acutely consuming the drug (Dourron et al., 2023). Other concerns include the potential for 5-MeO-DMT to induce seizures in individuals with a predisposition, or for it to increase body temperature to unsafe levels.

Our model predicts that a highly-biased 5-HT1a agonist such as 5-MeO-DMT will only have clinically relevant effects at very high doses, and this seems to be the case from early empirical studies (Reckweg et al., 2023). Increases in dose strength however produce decreases in psychological tolerability as well as corresponding undesirable side-effects resulting from high levels of 5-HT agonism at non-clinically relevant sites throughout the nervous system. An ideal balance between 5-HT2a and 5-HT1a agonism may involve a relatively smaller but pharmacologically meaningful bias towards 5-HT1 agonism over 5-HT2a agonism.

Notably, 5-HT itself is biased in this way with a three to five fold preference for 5-HT1a over 5-HT2a agonism (Barnes & Sharp, 1999), which is consistent with the optimal binding profile reported in our results. It may prove worthwhile to explore the development of drugs with a similar profile to 5-HT itself, but which are acutely psychoactive and amenable for use in psychedelic-assisted therapy. 5-MeO-DMT is part of a larger class of “5-methoxy” psychedelic substances which all posses greater binding affinity for 5-HT1a over 5-HT2a receptors (Puigseslloses et al., 2024). There is early evidence that this class of drugs may also posses a promising therapeutic profile (Warren et al., 2024), as would be predicted by our model.

### 4.4 Limitations and future directions

While important to affect and cognition, the receptor populations we consider here are only two of a dozen 5-HT receptor types expressed throughout the central nervous system. There is evidence that other 5-HT receptor populations including 5-HT1b and 5-HT2c may also play important roles in the etiology of certain mood disorders (Ruf & Bhagwagar, 2009; Millan, 2005; Carr & Lucki, 2011). These two populations in particular are also common targets of classic psychedelics including LSD and psilocybin, though they do not possess the same level level of binding affinity or causal influence as either 5-HT2a or 5-HT1a (Rickli et al., 2016; Deco et al., 2018). The action of these receptor populations on cortical activity is also less straightforward than the apparent opponency between 5-HT2a postsynaptic excitation and 5-HT1a postsynaptic inhibition. Further basic neuroscientific research will be required to fully understand the complex role that the family of 5-HT receptor populations plays throughout the cortex and the central nervous system more broadly.

Our theoretical model is based on a number of assumptions which are still in need of experimental validation. Our model operates on the basis of the Bayesian brain hypothesis, and assumes that cortical dynamics are governed by a process of PP (Keller & Mrsic-Flogel, 2018). More specifically, we make the assumption that PP can be modeled as learning and stochastic sampling in an energy-based model using gradient descent (Hoyer & Hyvärinen, 2002; Aitchison & Lengyel, 2016; Dong & Wu, 2023). Human behaviour is consistent with Bayesian inference in many sensory (Ernst & Banks, 2002), motor (Körding & Wolpert, 2004) and cognitive tasks (Tenenbaum et al., 2006), and while there is accumulating evidence for predictive processing in various aspects of cortical computation, it is still not universally accepted among theorists (Walsh, McGovern, Clark, & O’Connell, 2020). A similar state of evidence exists for explicit attractor dynamics in cortical computation, which would be amenable to modeling using energy functions (Khona & Fiete, 2022). While research into the existence of such dynamics is still nascent, there is accumulating evidence that within the context of high-level decision making PFC cortical dynamics can be modeled using energy functions (Mante et al., 2013; S. Wang et al., 2023).

Assuming that our theoretical assumptions are correct, our model still contains a few limitations which are worth discussing. First, we limit our consideration to the dynamics of only a single fixed target energy function and a single associated learned energy function. This is a significant simplification of real neural activity, where predictive processing is hypothesized to take place at many levels of a spatiotemporal hierarchy and likely involve non-linear feedback dynamics.

Second, we make the simplifying assumption that the environmental context which the organism finds itself in is held fixed for the duration of the simulation. Consequently, the generative model’s parameters, from latent neural responses to reconstructed stimuli, presumably context-dependent, are assumed fixed, and omitted throughout the drug experience. This allowed us to avoid considerations of how the external dynamics of the world may alter the sensitivity of the neural optimization process, particularly its response to sensory information, or attentional modulation of prediction errors over time. At the same time, it has prevented us from studying how the drug effect may impact lifelong or continual learning (Parisi, Kemker, Part, Kanan, & Wermter, 2018).

Third, we resorted to a tabular representation of the optimization landscape, which is modeled via a two-dimensional neural response **z**, which we update directly by gradient descent on the energy function. While this is a significant simplification of neural dynamics in living organisms, there is evidence to suggest that for a given functional network, the underlying optimization dynamics may often lay on a lower-dimensional manifold (Khona & Fiete, 2022). Indeed, multiple studies have found that cortical attractor dynamics can be projected down to as simple as a 1D space while retaining significant predictive power in the case of manifolds for high-level decision making and representation (Inagaki, Fontolan, Romani, & Svoboda, 2019; Okazawa, Hatch, Mancoo, Machens, & Kiani, 2021; S. Wang et al., 2023).

Fourth, our model does not take into account the role that action plays in the belief inference process. Given that perception and belief formation is ultimately in the service of adapted and goal-directed action, this would serve as a worthwhile extension to the model. The PFC in particular has been heavily studied for not only its role in high-level belief representation, but also for high-level action selection (Domenech & Koechlin, 2015; S. Wang et al., 2023). Numerous theoretical models currently exist which attempt to provide normative descriptions of this role of the PFC, often from the perspective of reinforcement learning, meta-learning, or both (J. X. Wang et al., 2018; Muller et al., 2024).

Fifth, it is the case that the simulations utilized in this work are abstracted from real neural data. Each of the starting energy functions are sampled from a set of structured noise functions, rather than being naturally induced by a specific set of parameters and perceptual or behavioral task. A promising future direction is to extend this work by considering datasets and excitatory-inhibitory models represented via recurrent neural networks which would themselves induce more ecologically valid energy functions. The currently expanding literature on work utilizing deep neural networks as models of brain function may be of particular interest towards realizing this goal (Aitchison & Lengyel, 2016; Barak, 2017; Richards et al., 2019; Juliani et al., 2024; Dong & Wu, 2023).

Finally, in order to realize the potential benefits of next-generation psychotherapeutic agents, we emphasize the importance of a measured and careful approach to any research in the mental health and psychedelic drugs space. Our model and simulation results are based on a number of theoretical assumptions which must ultimately be verified through further empirical research. It is also important that due attention be paid to issues of health equity and ethics to ensure that the consequences of such research are beneficial and equitably distributed. Considerations of individual difference in responses to serotonergic psychedelics is thus an important and under-explored area of research.

## 5 Conclusion

Psychedelic-assisted psychotherapy has the potential to improve the lives of many individuals suffering from cognitive, affective, and behavioral disorders which are not amenable to current treatment modalities (M. W. Johnson & Griffiths, 2017; M. W. Johnson et al., 2019). A critical step towards realizing this potential is to develop sophisticated models which can be used to predict the therapeutic efficacy of candidate psychedelic substances and doses. While the amount of clinical and neuroscientific evidence which can be used to develop such models continues to accumulate at a rapid pace (Carhart-Harris et al., 2021; Doss, de Wit, & Gallo, 2023), actionable models are still in their infancy.

One popular theoretical framework understands psychedelics primarily as pharmacological agents which engender greater neuroplasticity across various temporal scales (Olson, 2018; Calder & Hasler, 2023). While appealing for its simplicity, this model cannot account for the mediating role that the acute phenomenological experience and its correlated neural activity play in driving positive clinical outcomes (Yaden & Griffiths, 2020). In contrast, frameworks privileging changes to subjective experience have focused on the role of belief alteration (Safron, 2020; Letheby, 2021), typically in the form of belief relaxation as a result of psychedelic use both acutely and post-acutely (Carhart-Harris & Friston, 2019). Here we presented a theoretical model based on predictive processing and a set of simulations in which this hypothesized process of belief alteration can be formally studied. Our model is compatible with other recent dynamical systems models psychedelic activity (Hipólito et al., 2023), including “neural annealing theory” (M. Johnson, 2019; Gómez-Emilsson, 2021). We hope that it will be able to guide further clinical and neuroscientific research on this topic.

Within our framework, we modeled the acute action of psychedelics on belief representation as the result of two unique sets of effects, generated by the populations of 5-HT2a and 5-HT1a receptors present in the prefrontal cortex. We operationalize these effects as two different forms of modulation of the process which encodes and maintains belief representations. At a high level, we find that both the effects of stochastic perturbation through postsynaptic excitation (5-HT2a) as well as uniform smoothing through postsynaptic inhibition (5-HT1a) result in improvements in this optimization process across various metrics, thus pointing to a potential therapeutic effect for both. We characterize these effects in optimization terminology as 5-HT2a agonism introducing a transient overfitting of beliefs and 5-HT1a agonism introducing a transient underfitting of beliefs.

We predicts that mixed agonists which activate both 5-HT2a and 5-HT1a populations will have the most desirable properties, as they may balance both long-term therapeutic efficacy as well as acute drug tolerability. Furthermore, a bias towards preferential 5-HT1a agonism may provide an even greater relative benefit to tolerability with only a minimal decrease in long-term therapeutic efficacy. Interestingly, the psychedelics currently being most thoroughly investigated for their potential as clinical tools all have various levels of mixed 5-HT agonism. Rather than being an undesirable property to be engineered away, this mixed agonism may be key to the unique profile of subjective tolerability and therapeutic efficacy which psilocybin, LSD, and DMT seem to possess, and pure 5-HT2a agonists such as DOI and DOB may not.

Looking ahead, increasing clinical interest is being placed on the biased 5-HT1a agonist psychedelic 5-MeO-DMT (Dourron et al., 2023), a substance which is capable of inducing powerful and long-lasting antidepressant effects in minutes (Uthaug et al., 2019). Rather than focusing on the development of non-hallucinogenic psychedelics, which may prove to have limited efficacy outside of animal models (Yaden & Griffiths, 2020), significant evidence exists in support of exploring the pharmacological design space of psychedelic substances with a biased 5-HT1a binding profile similar to that of 5-MeO-DMT. Doing so may yield substances which are psychologically safer and more effective for clinical use than those currently under investigation. Such novel substances would enable the more mainstream adoption of psychedelic-assisted therapy, creating life-changing opportunities for the large population of individuals whose needs have been unmet by current front-line psychiatric treatments.

## 6 Acknowledgements

The authors thank Andrés Gómez Emilsson, Alex Meurice, Blake Richards, Colin Bredenberg, Michael Johnson, and Yad Konrad for the valuable insights they shared during preliminary discussions of the ideas eventually presented here. VC is grateful for support from Fonds de recherche du Québec – Nature et technologies (FRQNT), B2X (dossier 321675) and IVADO, Fonds d’excellence en recherche Apogée Canada, Bourse d’excellence au doctorat 2021.

## A Notation

**Table 1:** Notation.

|  |  |
| --- | --- |
| **Indices** |  |
| $t$ | outer-loop iteration index $t \in \{0, \dots, T - 1\}$ (neuromodulation) |
| $T$ | total number of outer-loop iterations. |
| $k$ | inner-loop iteration index $k \in \{0, \dots, k - 1\}$ (sampling) |
| $K$ | total number of inner-loop iterations (sampling). |
| **Parameters** |  |
| $\mathbf{z} \in \mathcal{Z}$ | neural response |
| $\mathbf{x} \in \mathcal{X}$ | stimuli |
| $\alpha$ | hyper-parameters of the neuromodulatory map/neuromodulatory strength/ learning-rates |
| **Maps** |  |
| $E^* : \mathcal{Z} \rightarrow \mathbb{R}$ | target energy function (ground truth) |
| $E : \mathcal{Z} \rightarrow \mathbb{R}$ | learning energy function |
| $\tilde{E} : \mathcal{Z} \rightarrow \mathbb{R}$ | auxiliary energy function for neuromodulation (exploration) |
| $\mathcal{L} \equiv D_{\text{KL}} : \mathcal{P}(\mathcal{Z}) \times \mathcal{P}(\mathcal{Z}) \rightarrow \mathbb{R}$ | KL-divergence between two distributions |

## B Theoretical and computational model

In this section, we give additional details concerning the theoretical framework in which the computation model operates and the theory surrounding it.

### Problem setting

Given stimulus **x** we seek a good neural representation, denoted with **z**, such that the neural response elicited encodes maximal information about the stimulus. Generally, this problem can be formalized as inference and learning in latent variable models and formally described as an optimization problem using the variational inference framework (Kingma & Welling, 2022). However in this work, we make a number of simplifications that allow us to bypass these methods. In particular, our problem can be defined in the following way. We assume oracle feedback about the optimal representation *p*^∗^(**z**) = exp(−*𝔼* ^∗^(**z**)) which is given as a target for the optimization procedure. The optimization problem is then to find the model *p*(**z**) ∝ exp(−𝔼(**z**)) that closely matches the optimal representation given the oracle feedback at each iteration. To fit an EBM to the target distribution we use the minimization objective

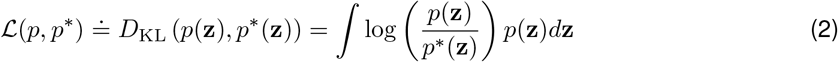

### Approximating the loss gradient

We want to differentiate and optimize the loss *ℒ* (*p, p*^∗^) with respect to the energy function *E*. Taking the functional gradient with respect to the energy function *E* yields

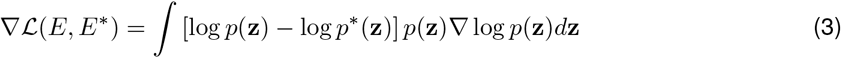

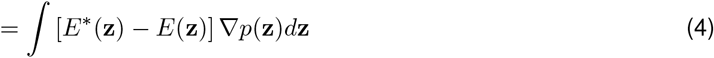

We make a number of approximations to estimate this gradient with respect to the energy function: (i) We generate samples from the (exploratory) neuromodulated model 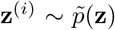, rather than *p*(*z*). (ii) We perform a semi-gradient operation in which we drop the terms of the gradient resulting from differentiating the partition function *Z*. (iii) We query the target oracle function for feedback *E*^∗^(**z**^(*i*)^). This yields the learning procedure

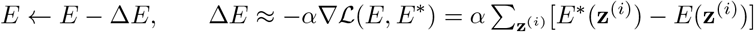

### Neural response sampling

We adopt a sampling method to generate neural responses from the EBM *p*(**z**). This means we can use Monte Carlo sampling (MCMC) to obtain the samples **z**^(*i*)^. There exist several MCMC samplers that can perform the procedure. The simplest of them is Langevin sampling, which uses

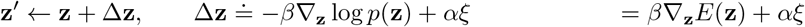

where the last equation follows from ∇_**z**_ log *p*(**z**) ∝ ∇_**z**_ log *p*(**z**) ∝ ∇_**z**_*E*(**z**).

A real concern with MCMC methods is that the Markov chain move through all areas of significant probability. If the target distribution has many modes or “islands” of high density, then it will take a long time to move from one island to another. When the initial proposal distribution used when sampling has a very large variance, then the chance of landing on a high-density island is small. Auxiliary variable MCMC methods such as stochastic Hamiltonian Monte-Carlo (HMC) and other neural adaptation procedures have been developed to address these concerns, making non-local jumps possible so that we can more easily jump from one mode to another. Our method uses samples from an exploratory neuromodulated model which allows us to bypass these issues. The idea is to use a auxiliary model, which accumulates the signal at each timestep *t* coming, e.g., from ion concentrations, release of neural transmitters, activation/inactivation of ion channels, drug-induced activation of receptors, etc.

This accumulation may be interpreted in different ways. One way to interpret it is simulating positive feedback loops within a recurrent network circuit which enhance the neural signal. Another is to view it as momentum or inertia that is being that acts to accelerate the dynamics, and thereby the sampling process, enhancing the exploration of the posterior.

## C Simulation Methods

The cortical dynamics simulator code was written in python, and is freely available to use with an open-source license at: https://github.com/awjuliani/serotonin-ebm. Table 2 presents the hyperparameters used in the experiments presented in this work. Section C.1 presents a set of studies which assess the sensitivity of our simulated results to values of a number of key hyper-parameters.

**Table 2:** Parameters used for simulation experiments.

| Parameter | Value |
| --- | --- |
| surface size ($N \times N$) | 50x50 |
| surface pattern | "noise" |
| energy function $\sigma$ | 5 |
| replications | 100 |
| total time-steps (T) | 400 |
| warm-up time-steps | 50 |
| drug time-steps | 200 |
| cooldown time-steps | 150 |
| beta distribution ($\alpha, \beta$) | (2, 5) |
| grad steps per time-step ($k$) | 10 |
| homeostatic $\alpha$ | 0.05 |
| plasticity $\alpha$ | 0.5 |
| dose level descriptors | [zero, min, light, medium, heavy, max] |
| Inhibitory $\alpha$ | [0.0, 0.03, 0.06, 0.13, 0.25, 0.5] |
| Excitatory $\alpha$ | [0.0, 0.03, 0.06, 0.13, 0.25, 0.5] |
| Inhibitory $\sigma$ | 2.5 |

### C.1 Simulator parameter sensitivity studies

We assessed the sensitivity of the simulated results to four key hyper-parameters; (1) Hebbian plasticity, (2) homeostatic strength, (3) the number of gradient steps taken when inferring the neural response (*k* in Algorithm 1), and (4) the energy function initializations.

#### Hebbian plasticity

Figure 6 presents results for a range of plasticity values. We find that our simulated results and overall conclusions remain consistent when plasticity *α* is varied from 0.1 to 1.0 (the default is 0.5). We do observe that when *α* = 1.0 the post acute number of local minima for mixed neuromodulators falls relative to the baseline plasticity value, however this does not meaningfully affect our conclusions as the difference is small relative to the acute changes in local minima count. However, when plasticity is very large (*α* = 2.0), our conclusions no longer hold. In this case divergence increases for all agonist mixes and strengths, including the baseline (zero). Here, we observe that the post acute gradient magnitude is substantially larger compared to the baseline plasticity, as is the number of local minima. This indicates the energy function has become more complex and it is easier to get stuck in sub-optimal local minima, likely leading to the observed increase in post-acute divergence. Note that since we observe the the baseline divergence increasing we do not think that 2.0 to be a very realistic value for plasticity *α*. We also present results for *α* = 0 to show the simulator dynamics in the absence of plasticity, however we note this is also not a realistic value for this parameter as it implies no learning is taking place.

**Figure 6.**
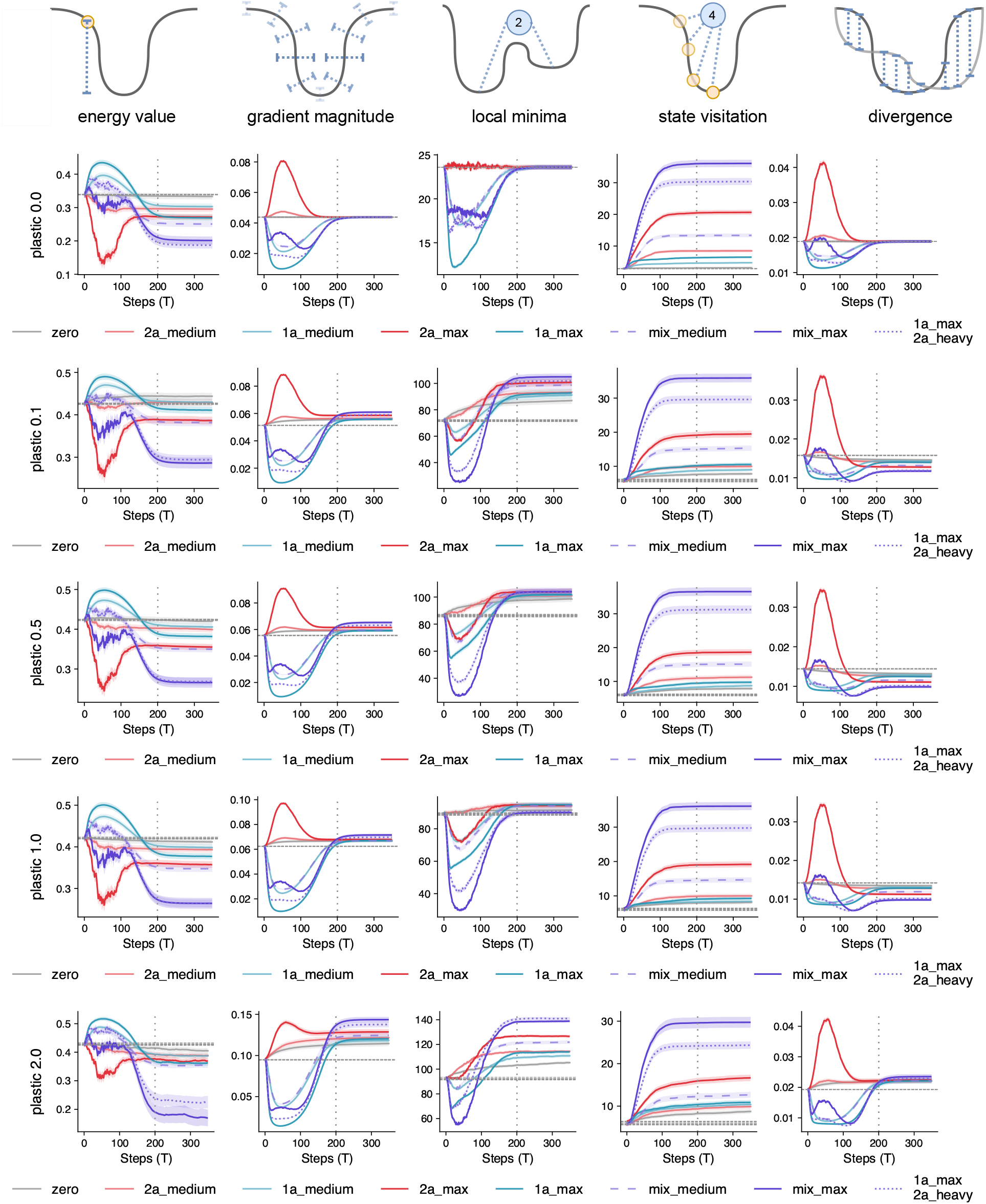
Hebbian plasticity hyper-parameter (*plastic*_*str*) sweep.

#### Homeostatic strength

Figure 7 presents results varying strengths of the homeostatic constraint. Overall we find that our simulated results and overall conclusions remain consistent when homeostatic *α* is varied from 0.025 to 0.1 (the default is 0.05). The weaker the homeostatic constraint, the more the acute drug phase effects are amplified. For example, compare the acute phase reduction in energy value or increase in gradient magnitude and divergence for 2a_max when *α* = 0.05 and *α* = 0.01. The stronger the homeostatic constraint, the faster the dynamics stabilize after the drug effect ends (vertical dotted line). When homeostatic *α* is very low, at 0.01, the energy value, gradient magnitude, local minima, and divergence all continue to change substantially for many time steps after the drug effect wears off. Further, the energy value and gradient magnitude have not stabilized by the end of the simulated experiment. This suggests that 0.01 is not a realistic value for this parameter. We also present results for *α* = 0 to show the simulator dynamics in the absence of any homeostatic constraint, however we note this is not a biologically plausible value for this parameter.

**Figure 7.**
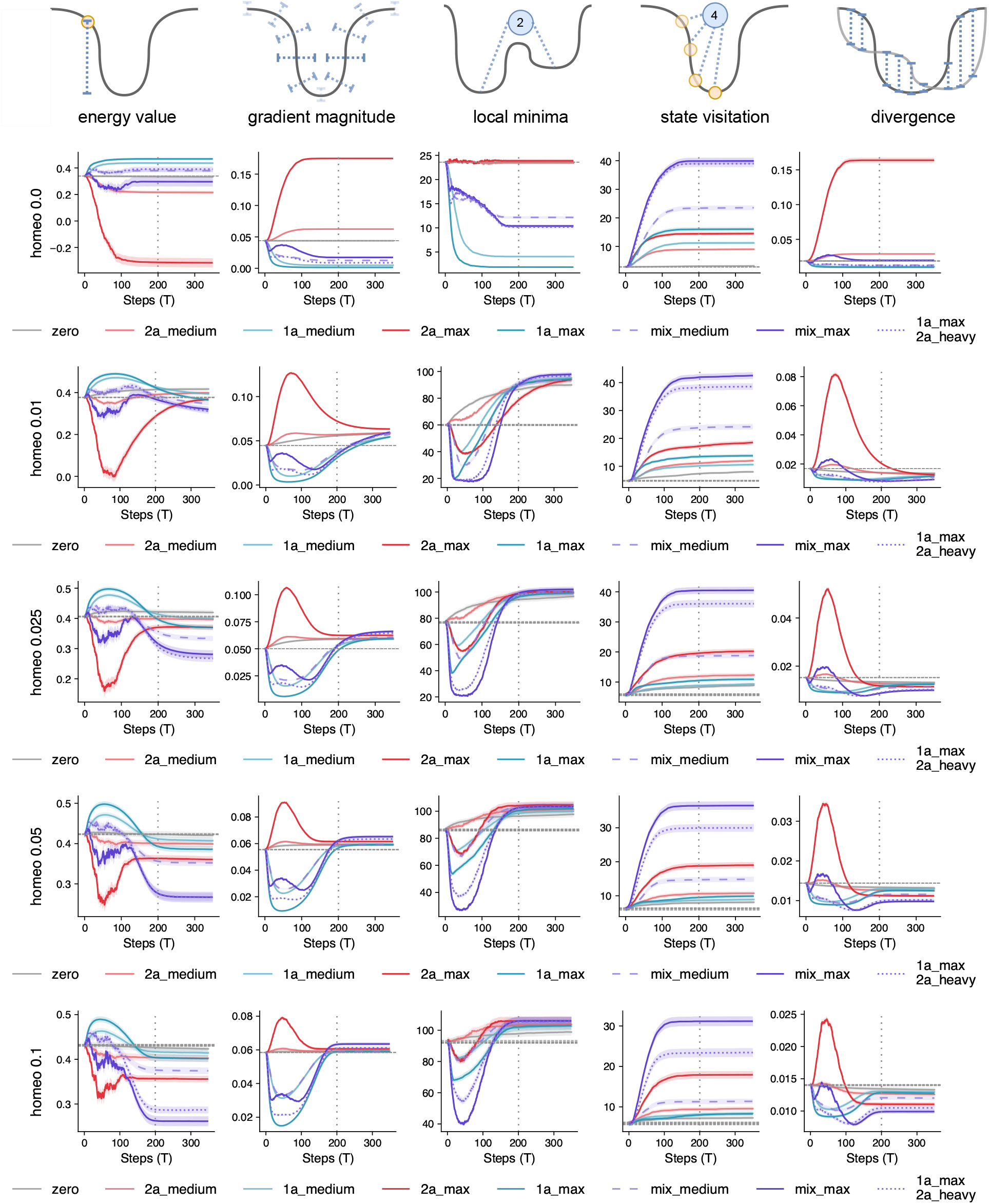
Homeostatic constraint hyper-parameter (*constraint*_*str*) sweep.

#### Gradient steps

We find that the simulated results are not particularly sensitive to the number of inference gradient steps each iteration, and that our overall conclusions would remain the same across a range of values. Figure 8 indicates that results are broadly consistent between 1 and 100 steps (the default is 10). Taking 100 or 10 gradient steps leads to almost identical results. When 1 gradient step is taken, the divergence and gradient magnitude is very similar to those when taking 10 or 100 steps. State visitation counts increase slightly while the number of local minima decreases marginally for the baseline and lower strength doses. The acute phase energy value for mixed modulators is somewhat sensitive to changes in the number of gradient steps. As gradient steps increase from 1 to 10, energy decreases during the acute phase, especially for the `mix_max` agonist. This is intuitive, since more gradient steps creates more opportunities for the neural response to descend the modulated energy surface.

**Figure 8.**
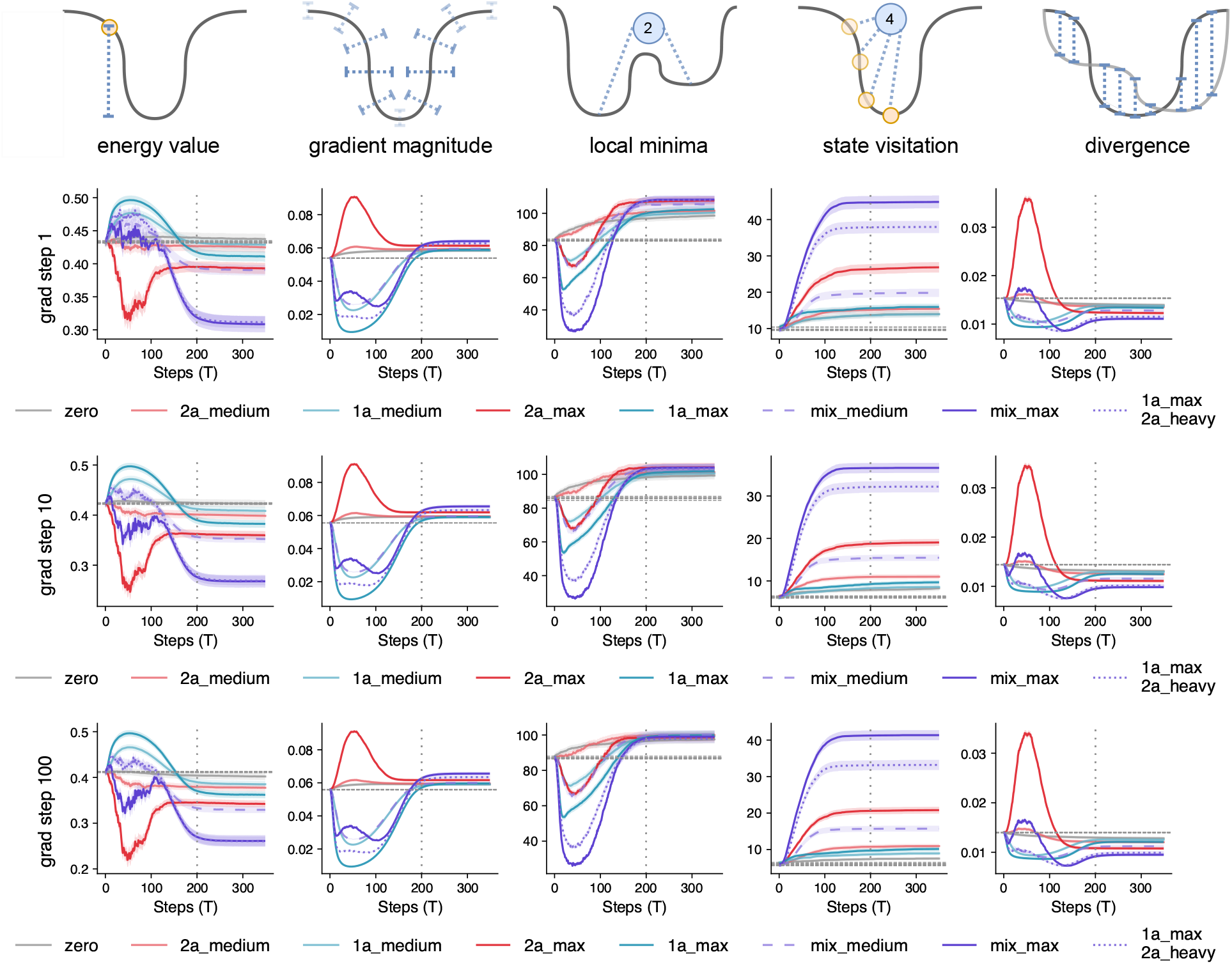
Number of gradient steps on z (*grad*_*step*_*num*) sweep.

#### Energy function initialization

We considered six different functions — `noise` (default), `log, sine, swirl, tangent, twisted sine` — for generating the initial energy function and the results are presented in Figure 9. Details of the functions are given in Table 3. We find that our main conclusions hold across a range different methods for initializing the energy functions *E*^∗^(**z**) and *E*_(_**z**). Additionally, in most cases the results and rank ordering of different modulators are consistent. There are two exceptions, the tangent landscape, and the energy value metric.

**Table 3:**
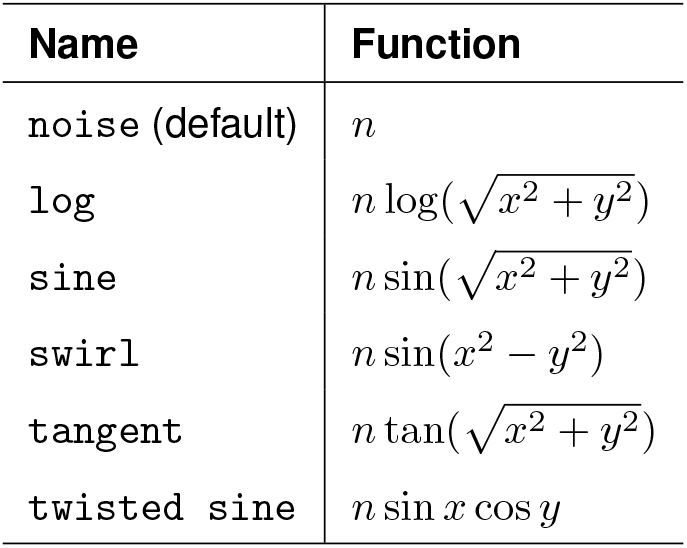
Initialization functions for *E*^∗^(**z**) and *E*_(_**z**). *n* ∼ *𝒩* (0, 1), (*x, y*) are coordinates of an *N × N* grid of points. Additional processing is performed by convolving the matrix with a guassian kernel *G*(*σ*) and normalizing the values to be within the range [0, 1].

| **Name** | **Function** |
| --- | --- |
| noise (default) | $n$ |
| log | $n \log(\sqrt{x^2 + y^2})$ |
| sine | $n \sin(\sqrt{x^2 + y^2})$ |
| swirl | $n \sin(x^2 - y^2)$ |
| tangent | $n \tan(\sqrt{x^2 + y^2})$ |
| twisted sine | $n \sin x \cos y$ |

**Figure 9.**
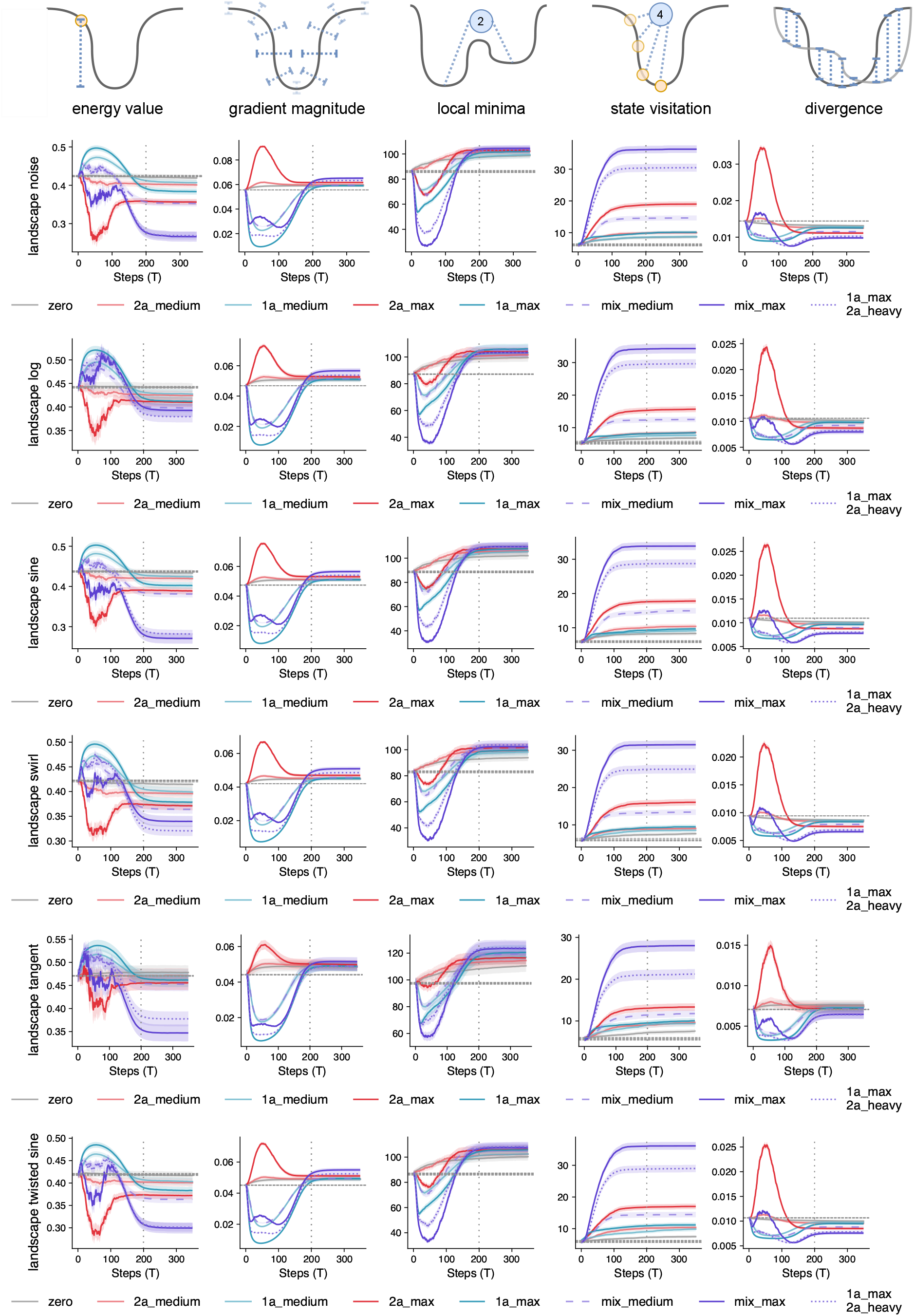
Energy function generator (*surface*_*pattern*) sweep.

When the tangent landscape is used, final KL-divergence values are higher and the error bars wider. However, consistent with our default generating function, we find that the mixed modulators `max-max, medium-medium` and `1a-max-2a-heavy` still result in the lowest final KL-divergence value and that the values are statistically significant [all *p <* 0.001] when compared with the baseline. Pure 2a agonism at max strength is also significantly lower than the baseline [*p <* 0.001]. Additionally, when we simulate all possible permutations of excitatory and inhibitory dose strengths (see Figure 10), the results are consistent with those presented in Figure 5. Interestingly, in this case a biased agonist of `1a-max-2a-heavy` has a relatively lower final KL-divergence value compared with the default noise landscape, and this value is close to the max-max combination. Finally, we note that the baseline divergence increases for this generating function, suggesting that the other choices of generating function are more suitable.

**Figure 10.**
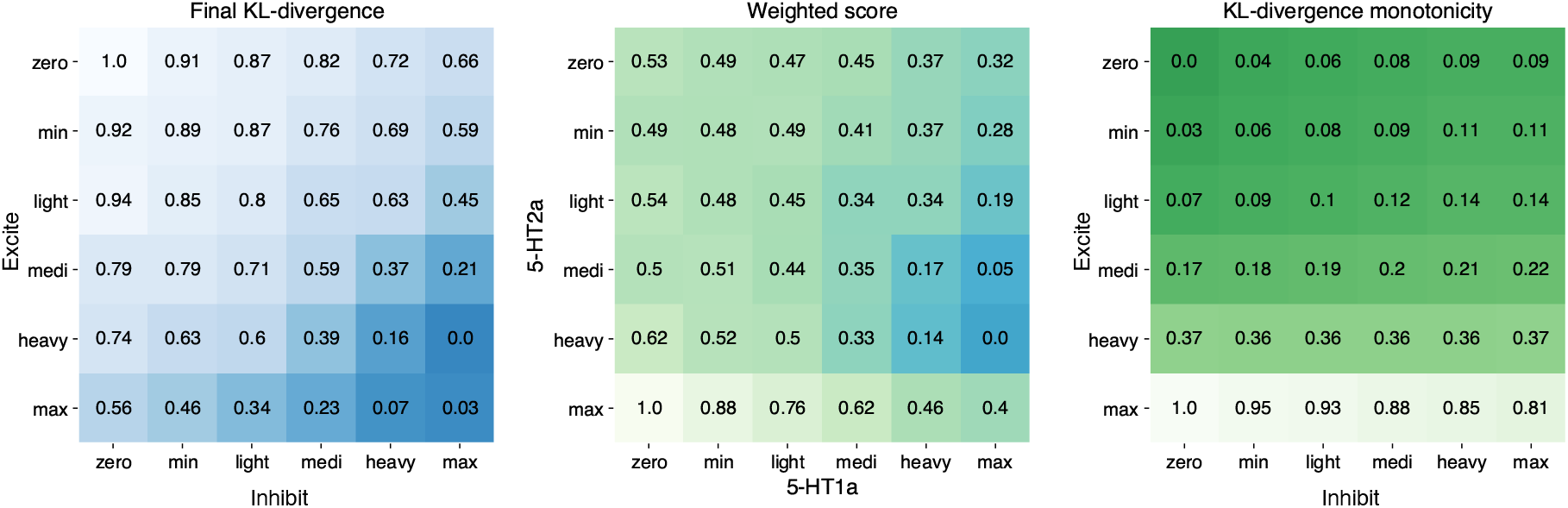
Matrix reporting normalized final energy values for set of possible excitatory and inhibitory modulation combinations using a tangent energy function initialization. The results and conclusions are consistent with those reported in Figure 5.

**Figure 11.**
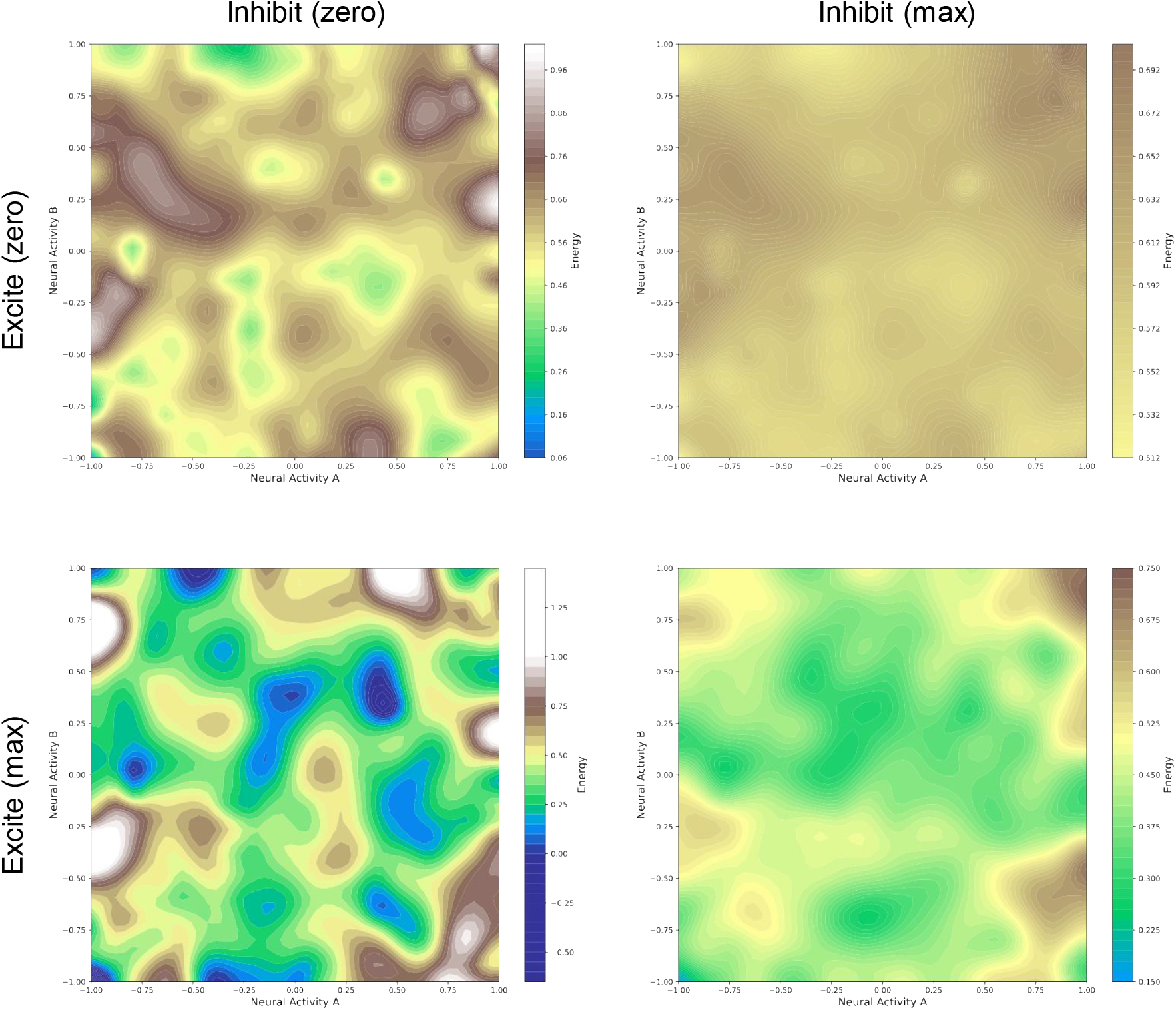
Representative examples of the energy function 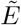 at the peak of the drug effect. **(Top Left)** No neuromodulation. **(Top Right)** Max level of inhibitory neuromodulation. **(Bottom Left)** Max level of excitatory neuromodulation. **(Bottom Right)** Max levels of both excitatory and inhibitory modulation.

Considering the energy value metric, we observe that the acute and post acute values for pure agonist (whether excitatory or inhibitory) are consistent across different generating functions. We consistently observe the acute phase transient over-fitting for excitatory modulators and acute phase transient underfitting for inhibitory modulators. However for mixed modulators, both the acute and post acute values appear sensitive to the choice of generating function. The acute phase may be characterized by an increase, decrease, or mixed change in the energy value. Additionally, the relative difference between the final energy value for mixed modulators compared to pure modulators, as well as the ordering between `mix-max` and `1a-max-2a-heavy` varies. What remains consistent across all landscapes is (1) that the final energy value for mixed modulators are lower than any acute phase value for the same agonist and (2) the final energy value of both the `mix-max` and `1a-max-2a-heavy` is lower than the baseline and any of the pure modulators. In 60% of landscapes (sine, tangent, twisted sine) the relative difference is similar to the baseline, in 20% of cases (swirl) the difference is moderately less but still substantial, and in 20% of cases (log) the gap is much smaller.

